# Genetic mapping of trait plasticity in a plant pathogenic fungus reveals genetic architecture and candidate genes for plasticity

**DOI:** 10.64898/2026.08.18.745209

**Authors:** Jessica Stapley, Bruce A. McDonald

## Abstract

Understanding how plant pathogens respond to environmental change is needed to better manage plant diseases. Phenotypic plasticity, the ability of a single genotype to produce different phenotypes across different environments, can influence pathogen adaptation and host-pathogen dynamics. Few studies have investigated the mechanisms underlying phenotypic plasticity in plant pathogens. Here we used phenotypic and genotypic data collected over >15 years and across multiple environments to perform genetic mapping of plasticity traits in the wheat pathogen *Zymoseptoria tritici*. Most (75%) of the QTL for plasticity (plQTL) overlapped with their corresponding mean QTL (mnQTL), suggesting that plasticity is controlled mainly by pleiotropic genes or tightly linked genes. 25% of the plQTL mapped to genomic locations separate from the mnQTL, suggesting that plasticity in these cases results from epistasis between unlinked loci. In several cases plasticity measured across different environmental gradients mapped to the same genomic positions, suggesting a shared control of plasticity for unrelated factors. These cases of shared control could be due to master regulators of plasticity or gene clusters. This mapping study provides unprecedented insights into the genetic architecture of plasticity in fungal plant pathogens.

## Introduction

Fungal plant pathogens pose a significant threat to food production globally. To counter this threat, significant resources are invested to control plant diseases, including deployments of fungicides, biological control agents, and disease resistant cultivars. A better understanding of how pathogen populations respond to changes in their environment and how this influences host-pathogen dynamics will enable better prediction and mitigation of the risks posed by pathogenic fungi to food production. In rapidly changing environments, phenotypic plasticity provides an important mechanism for plant pathogens to survive and adapt to new environments.

Phenotypic plasticity is defined as the ability of a single genotype to produce different phenotypes across different environments (Bradshaw, 1965; Via and Lande, 1985). Plasticity can be advantageous in changing or fluctuating environments and facilitate colonization of new environments, enabling a population to persist long enough for adaptive evolution to occur. This latter point is often referred to as the “plasticity-first” hypothesis (Levis and Pfennig, 2016). Plasticity can also facilitate adaptive evolution by releasing cryptic genetic variation (Paaby and Rockman, 2014). In the context of host-pathogen interactions, phenotypic plasticity can allow both hosts and pathogens to alter traits in response to environmental changes, affecting coevolution and disease dynamics (Taylor et al., 2006). Relatively few studies have investigated phenotypic plasticity in fungal plant pathogens or investigated the possible genetic control and genetic architecture of plasticity. A better understanding of the molecular mechanisms underlying plasticity can improve evolutionary models considering plasticity within host-pathogen interactions and better inform development of strategies to control existing and emerging pathogens.

Three models have been proposed to explain the genetic control and evolution of trait plasticity; overdominance, pleiotropy, and epistasis (Scheiner, 1993). The overdominance model, which is not relevant for haploid organisms, including many fungi, assumes that the variance of the phenotypes produced across environments by a multilocus genotype decreases as the number of heterozygous loci increases (Gillespie and Turelli, 1989). In the pleiotropic or allele sensitivity model, plasticity is caused by different alleles in the same gene having different sensitivity to variation in the environment, thus giving rise to variation in the phenotype. In simple terms, this model proposes that changes in the mean and variation in a trait in response to variation in the environment are determined by the same gene (Scheiner, 1993). In contrast, the epistasis or regulatory gene model posits that plasticity is caused by a separate, environmentally sensitive class of genes that regulate the expression of other genes that directly control the trait’s mean phenotype (Taylor et al., 2006). By simultaneously mapping a trait’s mean value and the plasticity for the same trait, we can differentiate between these models and determine if phenotypic plasticity is more often a result of pleiotropy or epistasis. The underlying genetic architecture influences how a trait can respond to selection (Kokko et al., 2017). If different genes or genomic regions control the mean value and the plasticity of the same trait, then the trait and its plasticity can evolve independently from one another.

There is little support for the overdominance model of plasticity (Pigliucci, 2005; Shende et al., 2024), while the pleiotropic and epistasis models are both well supported. For example, a study in *Caenorhabditis elegans* found co-localization of QTLs for body size plasticity and body size under different temperatures, supporting the pleiotropic/allele sensitivity model (Gutteling et al., 2007; Maulana et al., 2022). In contrast, genomic regions associated with thermal plasticity in body size in *Drosophila melanogaster* did not overlap with genomic regions associated with the corresponding trait mean – supporting the epistasis model (Lafuente et al., 2018). It is not surprising that some studies have found cases where both models are supported. In segregating *Saccharomyces cerevisiae* populations grown across 34 different environments, 72% of QTL associated with plasticity (plQTL) were associated with the trait mean (mnQTL) supporting the pleiotropy model, 6% of plQTL were not associated with the corresponding mnQTL, supporting the epistasis model, and the remaining 22% of plQTL were within 50 kb of the mnQTL, which was interpreted as further support for the pleiotropy model (Yadav et al., 2016). A study mapping QTL in barley in response to the presence of aphids and to variation in the rhizosphere found that 28% of the plQTL overlapped with the corresponding mnQTL (Tétard-Jones et al., 2011). A QTL study in tomato found 21% overlap between the trait mean and trait plasticity, suggesting that pleiotropy may be determining some plasticity, but for the majority of studied traits epistasis was considered more likely to control plasticity (Diouf et al., 2020). Recent studies in maize found a partial overlap in the genomic regions associated with both the trait mean and trait plasticity, supporting both the epistatic and pleiotropic models (Jin et al., 2023; Kusmec et al., 2017; Tibbs-Cortes et al., 2024).

The precision of mapping will clearly influence the interpretation of results from these studies. QTL intervals often contain many genes, but unless the causal mutation or gene for a trait is confirmed in follow-up functional studies, QTL studies cannot rule out the possibility that trait mean and trait plasticity are controlled by different but closely linked genes located in the same QTL interval. GWAS mapping studies generally provide more precise mapping unless there is a lot of linkage disequilibrium. One GWAS study in maize found that only four out of 977 SNPs were associated with both the trait mean and trait plasticity (Kusmec et al., 2017). Another GWAS study in maize that attempted to functionally annotate the significant intervals found a complex genetic architecture involving allelic heterogeneity, multiple alleles and pleiotropy (Jin et al., 2023). Thus an overlap between plQTL and mnQTL could be due to either pleiotropy in a single gene or the action of multiple tightly linked genes. Tightly linked sets of genes with similar functions or involved in the same pathway or biological process, often termed gene clusters or super genes, are common in many organisms. Within fungi the best characterized examples are biosynthetic gene clusters (BGCs), which can enhance adaptation to environmental stress through horizontal gene transfer, gene loss and rearrangements (Jia et al., 2021). Recent modelling has demonstrated that selection for plasticity can drive the evolution of recombination and the patterns of genetic distance between plasticity loci (Gulisija and Plotkin, 2017). Thus understanding the molecular mechanism controlling plasticity can provide insights into not only the relative roles of epistasis and pleiotropy in the evolution of plasticity, but also into the evolution of gene clustering and genome organization.

Although several studies have investigated the occurrence of quantitative phenotypic plasticity in fungi (Castiblanco et al., 2020; Kronholm et al., 2016; Muletz-Wolz et al., 2019; Pellon et al., 2022; Slepecky and Starmer, 2009), very few have mapped the genetic architecture of plasticity or identified genomic regions associated with trait plasticity. A large genetic mapping study with over 4000 segregants and 20 different media conducted in the yeast *Saccharomyces cerevisiae* found that plasticity is a polygenic trait, where 46% of QTL were shared between a trait’s mean and its plasticity (Zan and Carlborg, 2020). This study also found that epistatic networks were environmentally sensitive and reorganized following environmental change. Another study mapping growth plasticity in *S. cerevisiae* identified a gene encoding a sulfite pump (Ssup1) with strong pleiotropic effects, but did not explicitly investigate the overlap between mean and plasticity (Peltier et al., 2018). In the plant pathogen *Fusarium graminearum,* genetic mapping of plasticity was performed. Although this study did not specifically address the question of co-localization between mean and plasticity QTL, in the discussion it was noted that for one genomic region there was support for the epistasis model (Vajou, 2023). In the wheat pathogen *Zymoseptoria tritici*, the tolerance or sensitivity to environmental stress, which is a proxy for plasticity, was quantified across multiple environments (e.g. temperature gradients, presence or absence of salt, reactive oxygen species, and fungicide) and genetically mapped (Lendenmann et al., 2015; Stapley et al., 2025; Zhong et al., 2021). The tolerance or sensitivity measure was not considered a plasticity trait in the previous studies and those studies did not specifically address the question of co-localization. Instead, the previous studies focused on QTL intervals that did not overlap the mean trait value in the benign environment in order to identify candidate genes for a specific stress response. Evidence for pleiotropy was found for melanisation rate, growth rate and fungicide sensitivity, and this was attributed to variation in a polyketide synthase 1 (*Pks1*) gene cluster (Lendenmann et al., 2015). Further interrogation of this region and the candidate genes within the *Pks1* gene cluster revealed that differential regulation of the melanin regulator 1 gene (*Zmr1*) was responsible for variation in melanin accumulation, growth rate, and fungicide sensitivity (Krishnan et al., 2018).

Genetic mapping of trait plasticity provides excellent opportunities to understand the genetic architecture of plasticity and identify genes or genetic mechanisms that generate pleiotropy and epistasis. Possible molecular mechanisms for plasticity include transcription factors (TFs) and epigenetic changes that can alter gene expression. TFs sensitive to the environment can potentially alter expression in many genes, acting as master regulators of response to environmental perturbations and thus can be highly pleiotropic. Master regulators are important in morphological transitions in fungi, which are a discrete form of plasticity. In *Candida albicans* the switch between cellular states opaque-white influences virulence and is under the control of the TF master regulator *Wor1* (Mallick et al., 2016). Another well studied morphological switch is the yeast-hyphae switch in *C. albicans* and *Z. tritici*, which is triggered by environmental factors and accompanied by changes in a large number of genes, with mutants unable to transition between morphotypes having reduced pathogenicity (Chow et al., 2021; Francisco et al., 2023). In *Z. tritici*, key genes regulating this temperature sensitive transition have been identified, including the transcription factor *ZtMsr1* and the protein phosphatase *ZtYvh1* which regulate the yeast-hyphae transition (Francisco et al., 2023). The transition can be activated by high temperature, whereby heat stress induces intracellular osmotic stress that activates the cell wall integrity (CWI) and high-osmolarity glycerol (HOG) MAPK pathways. When cell wall integrity is compromised, ZtMsr1 inhibits hyphae formation (Francisco et al., 2023).

Highly connected loci, often called ‘hubs’, are loci connected within genetic networks through coordinated transcription/expression, often in close physical proximity and can be highly conserved and pleiotropic. An example is Heat Shock Protein 90 (HSP90). In *S. cerevisiae,* replacement of the wild type HSP90 with an orthologue from a species adapted to hyper-saline environments results in impaired growth in a low-saline environment and improved growth in a high-saline environment – demonstrating how variation in a hub locus can affect phenotypic plasticity (Kovuri et al., 2023). HSP90 also has the capacity to influence epigenetic mechanisms and change the chromatin states of many genes (Zabinsky et al., 2019). The role of epigenetic mechanisms in controlling phenotypic plasticity has been studied in the filamentous fungus *Neurospora crassa* (Kronholm et al., 2016). Kronholm et al (2016) showed that plasticity measured using reaction norms across four environmental parameters (temperature, osmotic stress, sucrose concentration and pH) varied across epigenetic knockout lines that differed in their presence of certain epigenetic mechanisms. The main finding of this work was that histone methylation and some small RNAs are likely to be the main epigenetic mechanisms involved in phenotypic plasticity (Kronholm et al., 2016). These examples illustrate how genetic studies of phenotypic plasticity can elucidate the relative roles of pleiotropy and epistasis in modulating plasticity and identify the genes and molecular mechanisms responsible for plasticity, including master regulators and/or epigenetic factors.

In this study we set out to genetically map phenotypic plasticity traits in the wheat pathogen *Z. tritici.* We used phenotypic data generated *in vitro* and *in planta* and genotypic data from experimental crosses collected over >15 years across multiple abiotic and biotic stressors (Table 1) to perform QTL mapping of plasticity. We also performed GWAS using phenotypic and genotypic data collected over >20 years across an international set of field populations sampled from five wheat fields on three continents.

**Table 1.**
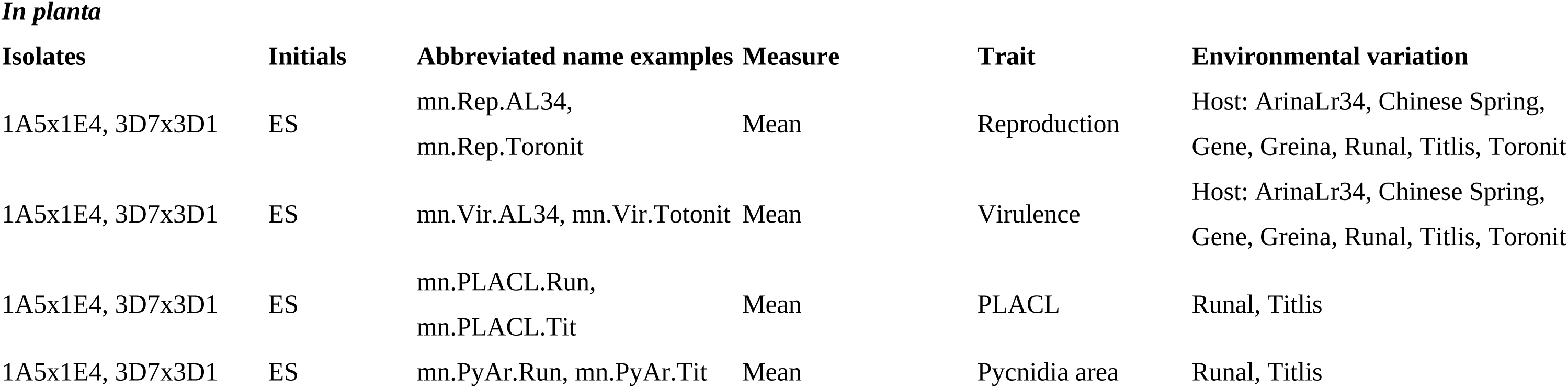
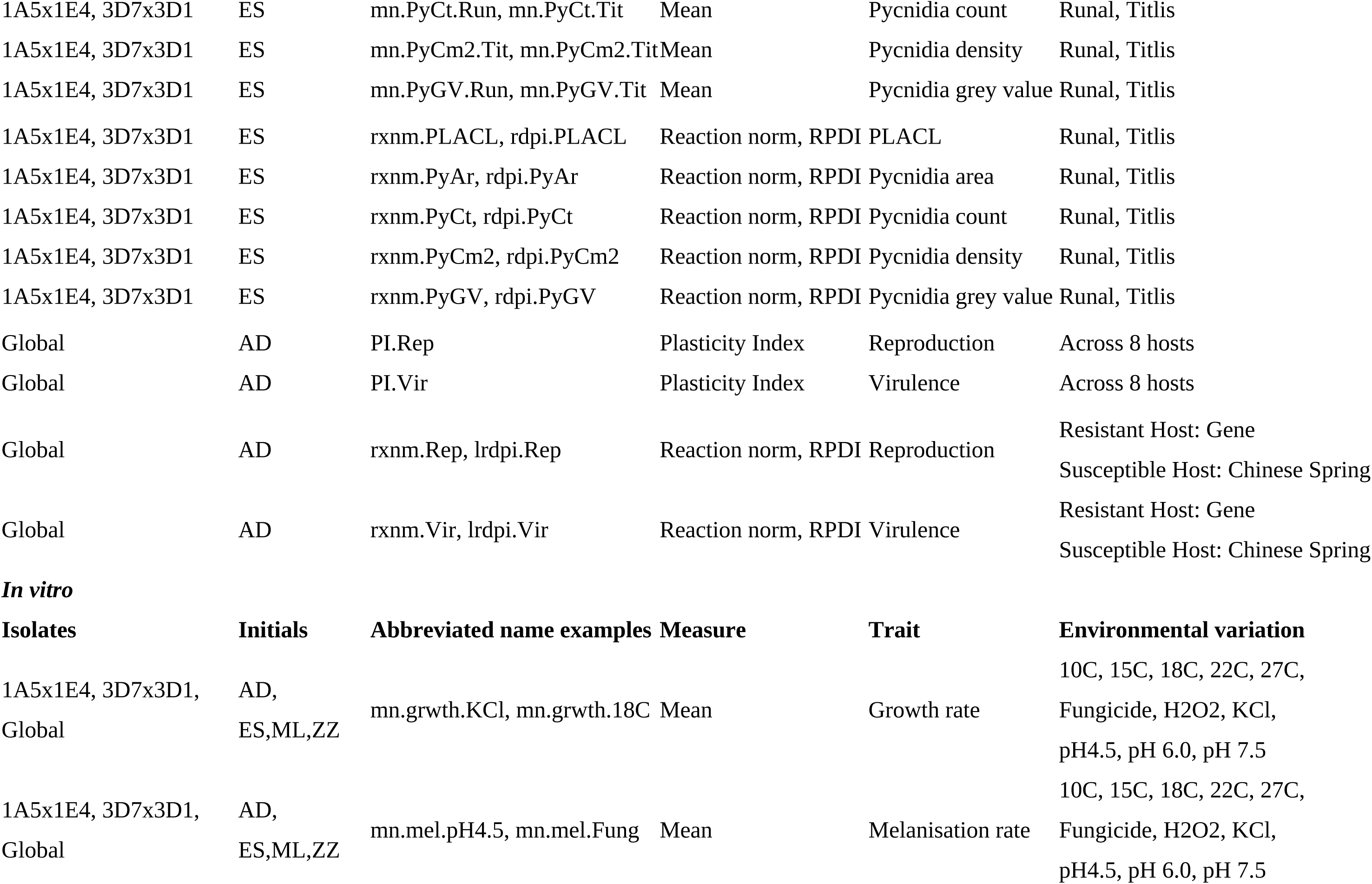
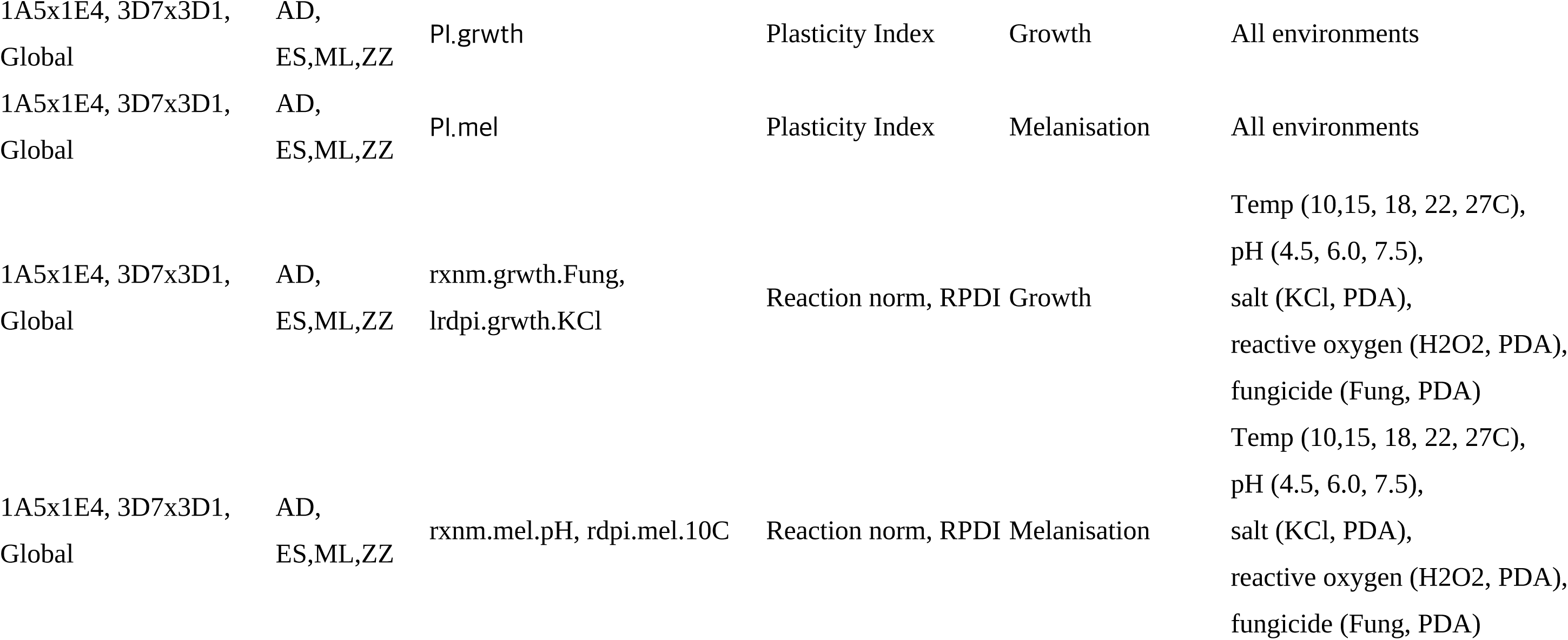
Overview of phenotypic data collected and key to abbreviated trait names. The data were obtained from offspring of two QTL populations (1A5x1E4 and 3D7x3D1) and a global population of strains by a primary data collector: Anik Dutta (AD) in 2019-2020, Ethan Stewart (ES) in 2010-2012, Mark Lendenmann (ML) in 2010-2012, and Ziming Zhong (ZZ) in 2015-2017. *In planta* traits were measured on different wheat genotypes, with Reproduction (Rep) (pycnidia density) and Virulence (Vir) (lesion area) recorded between 19-23 days post inoculation (dpi) and the percentage of leaf area covered by lesions (PLACL), pycnidia area (PyAr), pycnidia count (PyCt), pycnidia density (PyCm2) and pycnidia grey value (PyGV) measured at 23 dpi. Across *in vitro* experiments, two traits were recorded; the colony radial size and grey value, at two or three timepoints (AD: 8, 11, 18 dpi; ML: 8, 11, 14 dpi; ZZ: 8, 12, 15 dpi), which were converted to a growth rate and a melanisation rate. These *in vitro* traits were measured across variation in temperature and pH, and across variation in salt, hydrogen peroxide and fungicide concentration. Both the trait mean value and the trait plasticity were genetically mapped using QTL or GWAS. Trait plasticity was calculated in three ways: the reaction norm (rxnm), relative distance plasticity index (rdpi), and plasticity index (PI). The abbreviated trait names are formed by combining shorthand information for the type of trait (plasticity/mean), the trait measurement (growth/melanisation rate) and the environmental variation. A complete list of trait names and descriptions are provided in Supplementary Table S2.

We addressed four main research questions:

1. Can plasticity traits be mapped using QTL and GWAS approaches? If so, what is the genetic architecture (GA) of plasticity for these traits?
2. Does the GA of trait plasticity overlap with the GA for the mean of the corresponding trait?
3. Do plasticity measures across different environments map to the same or overlapping genomic intervals – representing putative plasticity hotspots?
4. Can we identify plausible candidate genes for trait plasticity?

Previously published QTL analyses of the *in vitro* data mapped the mean growth and melanisation rates and the sensitivity or tolerance in growth and melanisation rates in the control environment compared to a stress treatment ((e.g. KCl (Stapley and McDonald, 2023), fungicide (Lendenmann et al., 2015)). The sensitivity or tolerance measure was calculated as the ratio of the rate measured in the stress treatment and the rate measured in the control environment, where values >1 occur when the mean growth or melanisation rate was higher in the stress treatment compared to the control, indicating high tolerance to the stress. This tolerance measure can be viewed as a proxy for plasticity (Valladares et al., 2006). Thus we expected to find some overlap between previous results and the results of this study, however this study is novel for several reasons. First, we focus on trait plasticity, which we measure directly and more accurately utilising all the individual replicate data and not just means across replicates. Second, we quantify the overlap between the plQTL and mnQTL to address questions about the evolution of plasticity. In contrast to previous studies which focused on QTL intervals for traits measured in a stressful environment (H_2_O_2_, KCl, high and low temperatures) that did not overlap the QTL intervals for a trait measured in the control environment, here we focus on genes involved in the plasticity of the response to stress. Finally, this is the first time all of these data collected over a >20-year period have been analysed together within a shared experimental framework.

## Materials and Methods

We combined genetic and phenotypic datasets coming from multiple experiments carried out in the ETH Zürich Plant Pathology labs over a >20-year period to identify regions of the *Z. tritici* genome associated with phenotypic plasticity. The datasets encompassed two genetic mapping approaches: QTL mapping and GWAS. The phenotypic data included colony growth and melanisation rates measured *in vitro* in benign and stressful environments, as well as pathogen virulence and reproduction measured *in planta* across twelve wheat varieties.

Each group of *Z. tritici* strains was phenotyped in multiple *in vitro* environments, including a control environment of nutrient rich culture media kept at a fixed temperature (18^0^C or 22^0^C) that encompassed the experimentally-determined optimal growth temperatures of 18^0^C −25^0^C measured from more than 400 strains sampled from 28 wheat fields in 15 countries on five continents (Minana-Posada et al. 2025). The control environment was amended with sublethal concentrations of a fungicide (0.75 mg/L = 0.75 ppm propiconazole), reactive oxygen (H_2_O_2_ 1.0 mM) or salt (KCl 0.75 M), adjusted across a wide range of pH values (4.5, 6.0, 7.5) and grown at four additional temperatures (10, 15, 18, 22, 27^0^C). QTL analyses and/or GWAS analyses for all of these traits, except for the different pH values, have already been performed and published (Dutta et al., 2023, 2021; Lendenmann et al., 2016, 2015; Stapley et al., 2025; Stapley and McDonald, 2023; Zhong et al., 2021). However, in order to enable comparisons across experiments, the raw data had to be transformed as described below.

### Sources of *Z. tritici* strains used for genetic mapping

#### QTL mapping crosses

These crosses were described in detail previously (Lendenmann et al. 2014). In brief, two crosses were conducted between four Swiss strains, yielding a total of 700 offspring to use for QTL mapping. The cross between strains ST99CH1A5 and ST99CH1E4 (herein referred to as strains 1A5 and 1E4 respectively) produced 341 progeny. The cross between strains ST99CH3D7 and ST99CH3D1 (herein referred to as strains 3D7 and 3D1) produced 359 progeny. The original choice of strains to cross was based upon seven measured traits for each strain, including i*n planta* virulence, pycnidia size and density, and *in vitro* growth rates under different temperatures and in the presence of fungicides (Zhan et al., 2005). We used these datasets to calculate multiple phenotypic plasticity measures and perform QTL mapping.

#### GWAS strains

The strains used for GWAS mapping originated from four countries on three continents: Australia (n = 27), Switzerland (n = 32), Israel (n = 30) and United States (Oregon.R, n = 26; Oregon.S, n = 30) and were collected in 2001, 1999, 1991 and 1990, respectively ((Zhan et al., 2005); see Supplementary Table S1 for details). The two populations from Oregon were collected on the same day from two different wheat cultivars, Madsen (Oregon.R) and Stephens (Oregon.S), growing in the same field. All the other populations were sampled along transects from single, naturally infected wheat cultivars growing in the same field. We will refer to this population as the “global” collection of strains.

#### Phenotyping

Phenotype datasets came from experiments conducted by PhD students Mark Lendenmann (ML), Ethan Stewart (ES), Ziming Zhong (ZZ) and Anik Dutta (AD). Details of the experiments, the traits measured and the naming conventions are provided in Table 1. All traits are listed in the Supplementary Table S2. The protocols for strain recovery from −80^0^C storage, growth *in vitro* and the measurements of colony size and colony melanisation were described previously (Dutta et al., 2021; Lendenmann et al., 2014; Stewart and McDonald, 2014; Zhong et al., 2021). In brief, for *in vitro* phenotypes, Petri dishes containing potato dextrose agar (PDA) were inoculated with a spore solution (concentration 200 spores/ml) to generate several independent colonies on each Petri plate. Inoculated plates were maintained at 70% relative humidity (RH) and at different temperatures depending on the treatment. For *in planta* phenotypes, standardized spore solutions were applied to seedlings of different wheat cultivars grown under standardized greenhouse conditions conducive to infection. Infected leaves were harvested, flattened and high-resolution digital images were produced to accurately measure infected leaf area and numbers of fruiting bodies produced by each infection.

#### *In vitro* phenotyping

Across all *in vitro* experiments, two measurements (colony area and colony grey value) were recorded at multiple days post inoculation (dpi) using automated image analysis as described previously (Dutta et al., 2021; Lendenmann et al., 2014; Stewart and McDonald, 2014; Zhong et al., 2021). Colonies were grown on 3 or 5 replicate plates for each treatment. Measurements were taken at multiple time points which varied across experiments (Lendenmann: 8, 11, 14 dpi; Zhong: 8, 12, 15 dpi; Dutta: 8, 11, 14, 18 dpi), thus we standardized the data by dividing colony area and grey value by dpi. Colony grey value, a measure of melanisation, was measured on a scale of 0-255, where darker, more melanised colonies have lower values (0 = black, 255 = white). Standardised colony area was transformed to radial size by dividing it by π and taking the square root of this value. We used the standardised colony radius and grey value to calculate both growth rate and melanisation rate over time. These rates were calculated by taking the difference between the first and last measurements and dividing by the number of days in between. For melanisation rate, more melanised, darker colonies have lower grey values and thus the rate is negative. To make this rate easier to interpret in terms of increased melanisation, it was multiplied by −1 to make the rate values positive. The growth and melanisation rates were then used to calculate the mean trait values per strain and multiple estimates of trait plasticity as described below.

#### *In planta* phenotyping

The methods for strain regeneration and inoculation of plants were described in detail previously ((for the QTL populations (Stewart et al., 2018), for the GWAS experiment (Dutta et al., 2021)). In brief, the strains were recovered from long term storage and used to prepare concentrated spore suspensions in 100 ml flasks containing 50 ml of yeast sucrose broth (YSB). The strains were grown at 18^0^C on a shaker for 4-7 days. The resulting blastospores were collected and either stored as a frozen pellet or as a spore solution. Spore solutions of 5 × 10^6^ spores/ml were inoculated onto plants that were grown in the greenhouse (22:18^0^C (day:night), 70% RH, 16h or 18h photoperiod) after they developed a fully expanded second leaf. Between 19-26 dpi, the second leaf was removed and mounted onto A4 paper and imaged using a flatbed scanner (GWAS dataset) or a digital camera (QTL mapping dataset). The digital images were processed using automated image analyses described previously (Karisto et al., 2018; Stewart et al., 2018; Stewart and McDonald, 2014). The strains from the QTL populations were tested on two wheat varieties (Titlis and Runal) and the size, number, density, and grey value of the pycnidia were measured (pycnidia size, pycnidia count, pycnidia density (per cm^2^ leaf), pycnidia grey value), as well as the lesion area (percent leaf area covered by lesions, PLACL). The GWAS strains were tested on a panel of 12 wheat lines, which included five landraces (Chinese Spring, 1011, 1204, 4391 and 5254), six commercial varieties (Drifter, Gene, Greina, Runal, Titlis and Toronit) and a back-cross line (ArinaLr34). However, for our analysis we only used data from eight of these lines (Chinese Spring, Drifter, Gene, Greina, Runal, Titlis, Toronit and ArinaLr34), excluding data from four landraces (1011, 1204, 4391 and 5254) because the number of replications was lower for these four lines (median=3) compared to the rest (median=7), because of limited seed availability when the phenotyping was performed. For the GWAS strains the pycnidia density and lesion area were recorded and used as proxies for pathogen reproduction and virulence, respectively. In total there were seven *in planta* traits (five used for QTL, two for GWAS). For each trait we calculated the mean across replicates. For pathogen reproduction we log transformed the data and then took the mean. The plasticity measures are described below. All phenotypic data is provided in Supplementary Material hosted on the ETH public repository [ETH Repository DOI: XXX].

#### Quantifying Plasticity

Multiple measures of plasticity can be estimated from quantitative data (Valladares et al. 2006), with different measures capturing different aspects of plasticity (Zan and Carlborg, 2020). We used three measures of plasticity for this study – the reaction norm (RXN), the relative distance plasticity index (RDPI), and the plasticity index (PI) across all environments. These were chosen as they are commonly used in the literature and provide complementary estimates of plasticity for our datasets (Valladares et al. 2006; Zan and Carlborg 2020; Arenas et al. 2025). Another common measure, the Finlay–Wilkinson regression model, was not used because this is only suitable when a trait varies linearly with the environment, and for the temperature and pH environments we found non-linear relationships. Though Finlay-Wilkinson regression is a simpler method for assessing genotype stability, reaction norm provides a more comprehensive view of phenotypic plasticity and adaptation across environments.

### Reaction norm (RXNM)

Mixed effect models were used to calculate the reaction norm for each individual strain across environments - either pairwise: comparing benign and stressful environments, or across a range (>2) of differing environments. A random slope estimate was used as a measure of plasticity. In the case of normally distributed traits measured across two environments (control and treatment) a linear mixed effect model was used. For temperature and pH, measurements were made across >2 environments and a quadratic mixed effect model was used. For *in planta* virulence and reproduction in the GWAS dataset, we calculated the slope using only two wheat cultivars - the most resistant and the most susceptible wheat varieties (Gene and Chinese Spring respectively). As these reproduction and virulence datasets were zero-inflated we used a hurdle model (Beta, zero-inflated and hurdle Poisson) using the R package GLMMadaptive (Rizopoulos 2025). It was not possible to calculate a reaction norm for the *in planta* traits in the QTL populations (ES) because the replicate data was not available.

### Plasticity index (PI) = ((Maximum (mean of reps) – Minimum (mean of reps))/ Maximum (mean of reps)

This is an estimate of plasticity across multiple (≥2) environments, where no linear relationship is expected. The PI was divided by the number of environments, to control for the fact that not all strains were measured in every environment.

### Relative distance plasticity index (RDPI) = |mean(cntrl)-mean(trt)/mean(cntrl)| or | mean(susceptible)-mean(resistant)/mean(susceptible)|

This is an estimate of plasticity across the two most divergent environments.

### Phenotypic correlation analysis

Pairwise correlations between all phenotypic traits were calculated in R with the ‘*cor’* function and p-values were adjusted for multiple testing using Benjamini & Hochberg and Bonferroni methods implemented in ‘*p-adjust’* (R Core Team 2020). Comparisons of the absolute (ignoring the sign +/-) correlation coefficient (ACC) between traits for both the QTL and GWAS datasets was performed using the Wilcox test in R (*‘wilcox.test’)*. Heatmaps of ACC between all plasticity traits were created to identify possible clusters of traits that may be linked by a shared genetic control or make up a phenotypic module. Heatmaps were created using the R package klaR::corclust (Roever et al. 2004).

### Genotyping

#### QTL Genotyping

SNP genotype data for each set of progeny were obtained from a RAD sequence dataset previously produced in our lab ((first described in (Lendenmann et al. 2014)). In brief, the genome was cut using the restriction enzyme *Pst*l and the libraries were sequenced on an Illumina HiSeq2000 with paired-end sequencing. Complete genome sequences of the parental strains ((NCBI Biosamples: SRS383146 (ST99CH3D1), SRS383147 (ST99CH3D7), SRS383142 (ST99CH1A5), and SRS383143 (ST99CH1E4)) were used to SNP genotype the parents and offspring (Croll et al. 2013).

RADseq processing, variant discovery and linkage maps are described in detail at https://github.com/jessstapley/QTL-mapping-Z.-tritici and in an earlier publication (Stapley and McDonald 2023). In brief, RADseq reads were trimmed of adapters and low-quality sequences using trimmomatic (v0.35). The RADseq reads were mapped to a reference genome using bwa mem (v0.7.17). Reads from progeny of the 3D7x3D1 cross were mapped to the 3D7 reference genome and progeny from the 1A5x1E4 cross were mapped to the 1A5 reference genome. Variant calling was done using the GATK Germline Short Variant Discovery pipeline, following their Best Practice recommendations (https://gatk.broadinstitute.org/hc/en-us/sections/360007226651-Best-Practices-Workflows). After this we applied the following filters: only biallelic SNPs, parents had alternative alleles, <50% missing genotypes per marker, <50% missing genotypes per individual, mean read depth >3 and <30, depth quality (QD)>5, mapping quality >40, minor allele frequency >0.02 and <0.80, and SNPs in regions where the SNP density is >3 in 10 bp were removed. Linkage maps built using these SNPs were described previously (Stapley and McDonald 2023). After filtering and map building, 63181 SNPs were available for analysing the 3D7x3D1 population and 32806 SNPs were available for the 1A5x1E4 population. A summary of the two linkage maps is provided in Supplementary Material SM1 and the complete maps are available at (https://github.com/jessstapley/QTL-mapping-Z.-tritici).

### GWAS genotyping

The genome sequence datasets are described in detail elsewhere (Hartmann et al. 2018; Dutta et al. 2022). All raw data is available on the NCBI Sequence Read Archive (SRA), with sampling location, year and SRA accession numbers provided in Supplementary Table S1. In brief, sequences were trimmed, mapped to the *Z. tritici* reference genome IPO323 using Bowtie (v2.3.3) and duplicates were removed using MarkDuplicates in Picard tools (v.1.118). Variant calling was done with GATK (v.4.0.1.2) for single nucleotide polymorphism (SNP) calling and variant filtration. The GATK HaplotypeCaller was used on each isolate with the command -emitRefConfidence GVCF and -sample_ploidy 1 (*Z. tritici* is haploid). Joint variant calls were performed using GenotypeGVCFs on a merged gvcf variant file with the option -maxAltAlleles 2. We used VariantFiltration and SNPs were removed if any of the following filter conditions applied: QUAL < 250 (overall quality filter); QD < 20.0 (avoiding quality inflation in high-coverage regions); MQ < 30.0 (avoid calls from ambiguously mapped reads); −2 > BaseQRankSum > 2; −2 > MQRankSum > 2; −2 > ReadPosRankSum > 2; FS > 0.1. The final dataset was obtained by filtering with a genotyping call rate of ≥80% and for the GWAS we used a minor allele frequency cut-off of MAF=0.05. The final SNP dataset contained 716619 SNPs. Heritability of traits based on these SNP data was calculated using Bayesian regression models to estimate the Genomic Best Linear Unbiased Prediction (GBLUP) implemented in the R package ‘hibayes’ (Yin et al. 2025) (scripts: https://github.com/jessstapley).

### Genetic mapping

#### QTL mapping

QTL mapping was performed using the R (v 3.6.0) package ‘qtl2’ (v2_0.24) as described in detail on github (https://github.com/jessstapley/QTL-mapping-Z.-tritici). We scanned the genome for a single QTL per chromosome with the ‘*scan1*’ function using a linear mixed effect model. For continuous traits we fitted the kinship matrix in the model to control for the relatedness of individuals (i.e. we included a random polygenic effect). Models that take into account the genetic covariance between individuals can reduce the false discovery rate in QTL scans and outperform models that do not include this information in the model (Malosetti et al. 2011). The significance threshold for a QTL peak was determined by 1000 permutation tests and we calculated a Bayes Credible Interval (95%) to identify the interval size around the QTL peak.

#### GWAS analysis

The program GAPIT (Wang and Zhang 2021) and the model FarmCPU was used to perform the GWAS using 716619 SNPs. A total of three principal components were included in the model to account for the population structure. For the GWAS *in vitro* datasets the sample size (n) is 99, while for the *in planta* datasets n=143. We used the default p-value adjustment in GAPIT, which is the Benjamini-Hochberg FDR controlling procedure. GWAS mapping of the *in vitro* datasets did not yield any significant SNP associations, possibly due to the low sample size (n=99). Inspection of the Manhattan plots suggested a few loci were approaching significance, so we estimated a suggestive, less conservative p-value cutoff using a chromosome level Bonferroni p-value threshold (0.05 / number of SNPS across a chromosome) for the *in vitro* GWAS traits.

### Analysis of mapping results

We used the mapping results to: 1) investigate the genetic mapping and colocalization of QTL and GWAS SNPs, and; 2) identify candidate genes affecting traits and trait plasticity.

#### Identifying genomic regions harbouring QTL and GWAS SNPs associated with traits

The GWAS and QTL mapping approaches used different strain populations and different reference genomes (IPO323, 3D7, 1A5). In order to compare the mapping results across these different datasets and identify overlapping genomic intervals and genes within these intervals, we converted the QTL genomic positions from 3D7 and 1A5 to the corresponding genomic positions in IPO323. IPO323 is the most well annotated of the three genomes (Lapalu et al., 2025). To convert genomic positions in the 3D7 and 1A5 genomes, we first extracted the list of genes within a QTL interval using the annotation (gff) files of the corresponding reference genomes. Using this list, we then identified the orthologous genes and genomic coordinates in the IPO323 genome using data from an analysis performed across the genomes of 19 reference strains (Badet et al. 2020). Using IPO323 coordinates we could obtain a list of the most recent IPO323 genes and annotations within QTL intervals. The GWAS analysis was performed using the IPO323 genome, thus significant GWAS SNP positions were already in the IPO323 genome and we could directly obtain the list of genes with significant GWAS SNPs.

Using IPO323 genomic coordinates we could determine if QTL from different populations and experiments map to orthologous genic regions and if significant GWAS SNPs map to these intervals. To reduce false positives, we removed large intervals. Using the 90% quantile of QTL interval length as a cutoff led us to exclude 14 intervals that were larger than 1.50 Mb. We used custom R scripts to quantify the percent overlap between QTL by calculating the overlap length (in bp) of a focal QTL with all other trait QTL intervals and dividing the overlap length by the focal QTL interval length. To investigate the colocalization of plasticity traits and mean traits, including traits measured across experiments and different environmental gradients (e.g. temperature, salt, fungicide stress), we collapsed overlapping QTL intervals and GWAS SNPs into distinct genomic regions (GRs). We then investigated the traits mapping to these genomic regions to address five questions (specified below) that are important for understanding the evolution and genetic control of phenotypic plasticity.

#### Q1) Do plasticity QTL or GWAS loci overlap with QTL or GWAS loci for corresponding mean of the trait?

How distinct are the genetic architectures or genetic loci affecting trait plasticity and trait mean values? If a trait’s plasticity is governed by pleiotropy, we expect that the mean trait value and plasticity for that trait will be controlled by the same genes and thus map to the same regions. If a trait’s plasticity is governed by epistasis, we expect that the trait mean value and plasticity for that trait will be controlled by different genes and thus map to different genomic regions. To investigate the overlap in genetic loci (i.e. SNPs used in either QTL or GWAS analyses) associated with the plasticity and the mean of the trait, we only considered the plasticity trait and its corresponding mean(s), specifically the plasticity and mean must be from the same type of trait (e.g. growth, melanisation, or pycnidia count), measured in the same population (1A5x1E4, 3D7x3D1), and measured in the same environments. For example, for RXNM in growth rate across temperature variation, there are five corresponding means; mean growth rate measured at 10, 15, 18, 22 and 27^0^C. For plasticity index the mean of all the traits measured in the population are the corresponding means. For example, for the PI for *in planta* reproduction there are eight corresponding means, one for each host.

#### Q2) Is there overlap between the SNPs identified in the QTL and GWAS mapping populations?

We counted the number of GWAS SNPs that map to a genomic interval. Addressing this question allows us to evaluate the consistency between different genetic mapping populations and methods.

**Q3) Is there overlap between QTL intervals across *in vitro* and *in planta* experiments (ZZ/ML and ES**)**?**

We counted the number of genomic regions that habour QTL for *in vitro* and *in planta* traits collected in the same mapping populations. If *in planta* and *in vitro* traits are mapping to the same regions, this suggests some shared genetic control of traits that may have very different roles in the pathogen’s life history.

#### Q4) Is there overlap between the experimental populations (1A5x1E4 and 3D7x3D1)?

We counted the number of QTL intervals for traits collected during different experiments (conducted at different times) and on different populations. We considered both mean and plasticity traits. Previous studies identified QTL for mean traits in these two populations, however with the exception of Lendenmann (Lendenmann et al. 2014; Lendenmann et al. 2015; Lendenmann et al. 2016), they did not search for overlaps between populations because they used different reference genomes for each population (Zhong et al. 2021; Stapley et al. 2025).

#### Q5) Is there overlap among QTL for plasticity measured across environments?

Considering only QTL for plasticity traits, we quantified how often the QTL for plasticity measured in one environment overlapped with QTL for traits measured in a different environment. For example, we compared the plasticity QTL found for variation in temperature with the plasticity QTL found for variation in KCl. When plasticity loci are shared between different environments it suggests that plasticity may be controlled by master regulatory genes that operate across a broad range of environments as opposed to specific regulatory genes that respond to a limited number of environments.

### Identifying candidate genes for plasticity within mapped genomic regions

**(a) GO term enrichment analysis.** We combined all genes found within the mapped plasticity QTL genomic regions to create a list of ‘genes of interest’. GO enrichment analysis was then performed following a tutorial for non-model species (https://archetypalecology.wordpress.com/2021/01/27/how-to-perform-kegg-and-go-enrichment-analysis-of-non-model-species-using-r/). GO annotations were retrieved from the annotation file for IPO323 (5593/13414 genes with GO terms) and we used the R package ‘topGo’ (Alexa and Rahnenfuhrer 2023) to perform a Fischer test on the genes of interest following the associated guidelines. All three ontologies – Biological Process (BP), Cellular Component (CC) and Molecular Function (MF) were analysed. The background set for GO analysis was the entire gene/transcript set for the reference IPO323 genome. KEGG pathway analysis was not included in the Results because K numbers can be found for only 4257 genes (31.7%) and the analysis yielded no enriched terms.
**(b) Biosynthetic gene cluster (BGC) enrichment.** To test whether each genomic region (GR) contains more biosynthetic gene clusters (BGCs) than expected by chance, we used a simulation-based approach. We used 34 BGCs identified in silico in *Z. tritici* (Cairns and Meyer, 2017). For this analysis we excluded intervals containing only GWAS-SNPs as these were too small. We simulated 1000 random genomic intervals on chromosomes 1-16, with the interval width drawn from a log-normal distribution approximating the real GR widths. We tested if the number of overlapping intervals across all GRs, and the number of BGCs within individual GRs were different to the equivalent numbers from the random intervals, providing an overall enrichment measure as well as enrichment measures at the level of individual GRs. An empirical p-value was calculated as the proportion of simulated intervals containing at least as many BGCs as the observed GR. All simulations were performed in R using the GenomicRanges package.
**(c) Focus on small intervals with high LOD scores**. To identify the most promising candidate genes, we focused on narrow genomic regions with high LOD (>10) or strong GWAS associations (high (>10) negative Log p-value). This method was used successfully in past investigations of *Z. tritici* to identify candidate genes affecting virulence (Zhong et al. 2017), morphology (Francisco et al. 2023) and melanization (Krishnan et al. 2018) that were functionally validated.

### Coding sequence comparison across parents

The *Hex-1* and *HSF* coding sequence was compared across IPO323 and the four parent strain reference genomes using BLAST+ (v2.17.0) and the analysis code was written with AI assistance (ClaudeOpus 5, ‘calude-opus-5’ Anthropic). Analysis details are in the github repository (https://github.com/jessstapley/QTL-mapping-Z.-tritici/). The AI alignments were critically evaluated by repeating the alignment in Aliview (v1.27) and verified by the authors to ensure accuracy.

## Results

In total we investigated 110 traits (54 mean, 56 plasticity). A total of 72 were phenotyped in the two progeny populations (1A5x1E4, 3D7x3D1) and mapped using QTL: 62 of these traits were measured *in vitro*, 10 traits were measured *in planta*. The other 38 traits were measured in the global population and mapped using GWAS: 22 of these traits were measured *in vitro*, 16 were measured *in planta*.

### Correlations of plasticity measures with each other and their corresponding means

All pairwise correlation coefficients for each population are provided in Supplementary Material SM2-SM5). Across all traits we tested if the plasticity trait and the corresponding mean trait had higher absolute correlation coefficient (ACC) when compared to all possible pairs of traits. A plasticity trait and the corresponding mean must be the same type of trait (e.g. growth, melanisation, or pycnidia count), measured in the same experimental population and across the same environmental variation. For example, for RXNM in melanisation rate across variation in KCl measured in 1A5x1E4, there are two corresponding mean traits: the mean melanisation rate at 18^0^C (control) and the mean melanisation rate in KCl measured in 1A5x1E4. In the QTL populations, the ACC between the plasticity trait and its corresponding mean were higher than ACC between all traits (Supplementary Figure S1). For the 1A5x1E4 population: Wilcox test statistic (*W*)=91672, *p*<0.001, mean ACC (mACC) across all pairs=0.13, mACC between plasticity and mean=0.37. For the 3D7x3D1 population: *W*=104090, *p*<0.001, mACC across all pairs=0.14, mACC between plasticity and mean=0.34. In the global population, a different pattern was observed. For *in planta* and *in vitro* traits the ACC between plasticity and the corresponding mean was not different to the ACC for all pairs: *W*=4140, *p*=0.43, mACC across all pairs=0.27, mACC between plasticity and mean=0.29; for *in vitro* traits: *W*=1344, *p*=0.20, mACC across all pairs=0.26, mACC between plasticity and mean =0.37.

Heatmaps of ACC for the plasticity traits showed a different clustering pattern across the different datasets (Supplementary Figure S2). In the 1A5x1E4 population there was one large cluster of 13 plasticity traits measured *in vitro*, these included plasticity in growth and melanisation rates across variation in salt and reactive oxygen concentration, temperature extremes and low pH (lrdpi.grwth.KCl, rxnm.grwth.KCl, lrdpi.grwth.H2O2, rxnm.grwth.H2O2, rdpi.grwth.10C, rdpi.mel.27C, lrdpi.mel.pH6.0, lrdpi.mel.pH4.5, lrdpi.mel.KCl, lrdpi.mel.H2O2, rdpi.mel.10C, rxnm.mel.KCl, rxnm.mel.H2O2). In addition, three *in planta* traits are correlated with each other forming a small cluster (rdpi.PyCm2, rxnm.PLACL, rxnm.PyCm2). In the 3D7x3D1 population there were three clusters of correlated traits, one involving plasticity of *in vitro* melanisation rate across nine traits involving variation in salt, reactive oxygen and fungicide concentration, temperature and pH (rdpi.mel.15C, lrdpi.mel.Fung, rxnm.mel.Fung, lrdpi.mel.pH6.0, lrdpi.mel.KCl, lrdpi.mel.H2O2, rxnm.mel.Temp, rxnm.mel.H2O2, rxnm.mel.KCl). The second group contained plasticity for eight *in planta* traits (rdpi.PLACL, rxnm.PLACL, rdpi.PyGV, rxnm.PyGV, rdpi.PyCm2, rdpi.PyCt, rxnm.PyCm2, rxnm.PyCt) but did not include pycnidia grey value. The third cluster was composed of plasticity for five *in vitro* growth rate traits involving variation in salt and reactive oxygen concentration and temperature and pH (lrdpi.grwth.pH7.5, rdpi.grwth.10C, rxnm.grwth.Temp, rxnm.grwth.H2O2, rxnm.grwth.KCl). In the GWAS dataset there are two groups of correlated *in vitro* traits. One group includes five traits involving growth rate or melanisation in the presence of fungicide or across temperature variation (rdpi.mel.15C, rxnm.grwth.Temp, PI.growth, rxnm.grwth.Fung, rdpi.mel.Fung). The other group of five traits includes plasticity in melanisation rate across all environments (PI), across variation in temperature and fungicide concentration and plasticity in growth rate across variation in temperature and fungicide concentration (rdpi.grwth.Fung, PI.mel, rxnm.mel.Fung, rxnm.mel.Temp, rdpi.grwth.15C). The three different plasticity measures for virulence are also clustered and the same pattern is found for reproduction. It is notable that plasticity in growth rate and melanisation rate group together in the global population and the1A5x1E4 population, whereas they group separately in the 3D7x3D1 population, with one group for plasticity in growth rate traits and a separate group for melanisation rate traits.

### Heritability

Heritability (h^2^) of plasticity was lower than the heritability of mean traits across all datasets (*F_1,180_*=11.72, *p*<0.001, for trait plasticity: mean h^2^=0.21, for trait mean: mean h^2^=0.28). For all traits, heritability was similar across the datasets, with a few exceptions (Supplementary Figure S3), namely *in planta* mean and plasticity measurements from the 1A5x1E4 population and *in vitro* plasticity measurements from the 3D7x3D1 population were lower than the heritability of mean *in vitro* measurements from the 3D7x3D1 population (*F_11,170_*=4.64, *p<*0.001).

### Genetic mapping and colocalization

From the 111 (QTL: 73, GWAS: 38) traits that were mapped, we identified 177 significant associations encompassing 141 QTL intervals and 36 GWAS SNP associations, spread across 14 chromosomes (Chr) (Chr1-12,14,16) Table 2, Figure 1). Four chromosomes had ≥20 QTL (Chr8 nQTL=32, Chr3 nQTL=24, Chr11 nQTL=22 and Chr7 nQTL=21). No significant QTL or GWAS associations were found for 44 traits (Supplementary Table S3). As expected for quantitative traits, there was a positive relationship between chromosome length and the number of QTL or GWAS associations (Pearson correlation coefficient (Pcc)=0.60, *t*=3.35*, p*=0.003, df=19) (Supplementary Figure S4). However, if we consider only the 14 core chromosomes there is no relationship (Pcc=0.10, *t*=0.34*, p*=0.73, df=10). Considering only the plasticity traits, there were 85 QTL or GWAS SNPs distributed across 11 chromosomes (Chr1-8, 10-12), with most mapping to Chr8 (18), followed by Chr7 with 11 and Chr3 with 10 mapped plasticity traits. Similar to what we found for all traits, there was a positive relationship between chromosome length and the number of QTL or GWAS associations (Pcc=0.63, *t*=3.57*, p*=0.002, df=19), but this correlation disappeared if we considered only the core chromosomes (Pcc=0.10, *t*=0.34*, p*=0.73, df=10).

**Figure 1.**
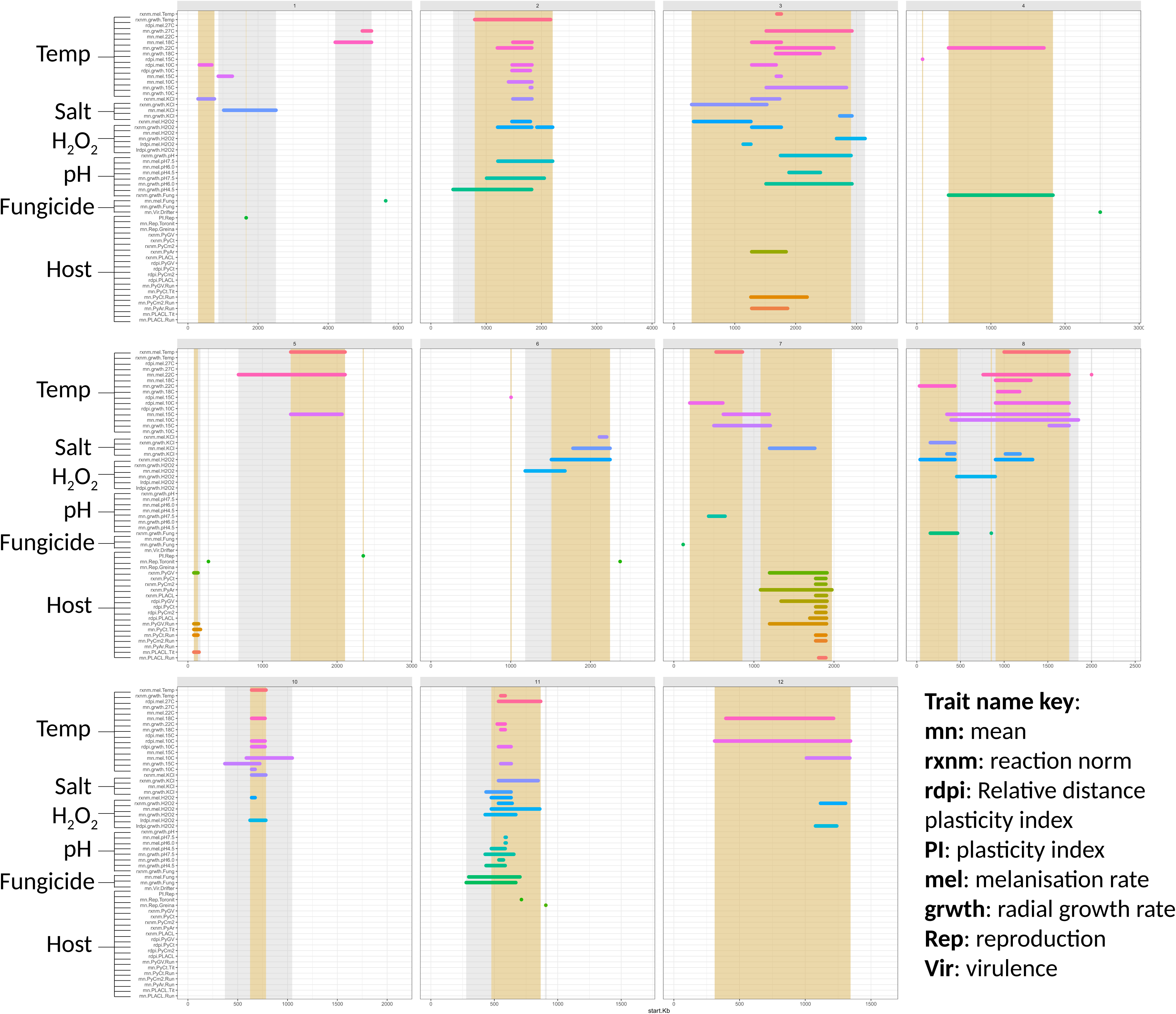
The chromosome-scale distribution and overlap of all significant QTL (intervals) and GWAS (SNP) associations for *in vitro* traits (growth rate and melanisation rate) across variation in temperature (Temp: 10, 15, 18, 22, 27^0^C), KCl concentration (Salt), hydrogen peroxide concentration (H_2_O_2_), pH (4.5, 6.0, 7.5), fungicide concentration (propiconazole), and *in planta* traits (Host) measured in two wheat genotypes (host: Titlis (Tit) or Runal (Run); traits: PyGV: pycnidia grey value; PyCt: Pycnidia count; PyCm2: Pycnidia per cm2; PyAr: Pycnidia area, PLACL: Percent leaf area covered by lesions), or measured in eight wheat genotypes (Host: Chinese Spring, Drifter, Gene, Greina, Runal, Titlis, Toronit and ArinaLr34), Vir: virulence, Rep: reproduction). Traits include three plasticity measures (PI, rdpi, rxnm) and mean (mn) measures. Numbers on top of each panel refer to chromosome number. Numbers along the bottom of each panel refer to base pair positions along each chromosome. Shading indicates the genomic regions where overlapping associations (QTL/GWAS) have been grouped, grey shading is shown for all trait associations, yellow-brown is shown for trait plasticity only. The colour-hue of the lines is a visual aid that links traits measured in different environments, pink-purple: temperature, blue: KCl and H_2_0_2_, aqua: pH and fungicide, green-brown-orange: *in planta*.

**Table 2.**
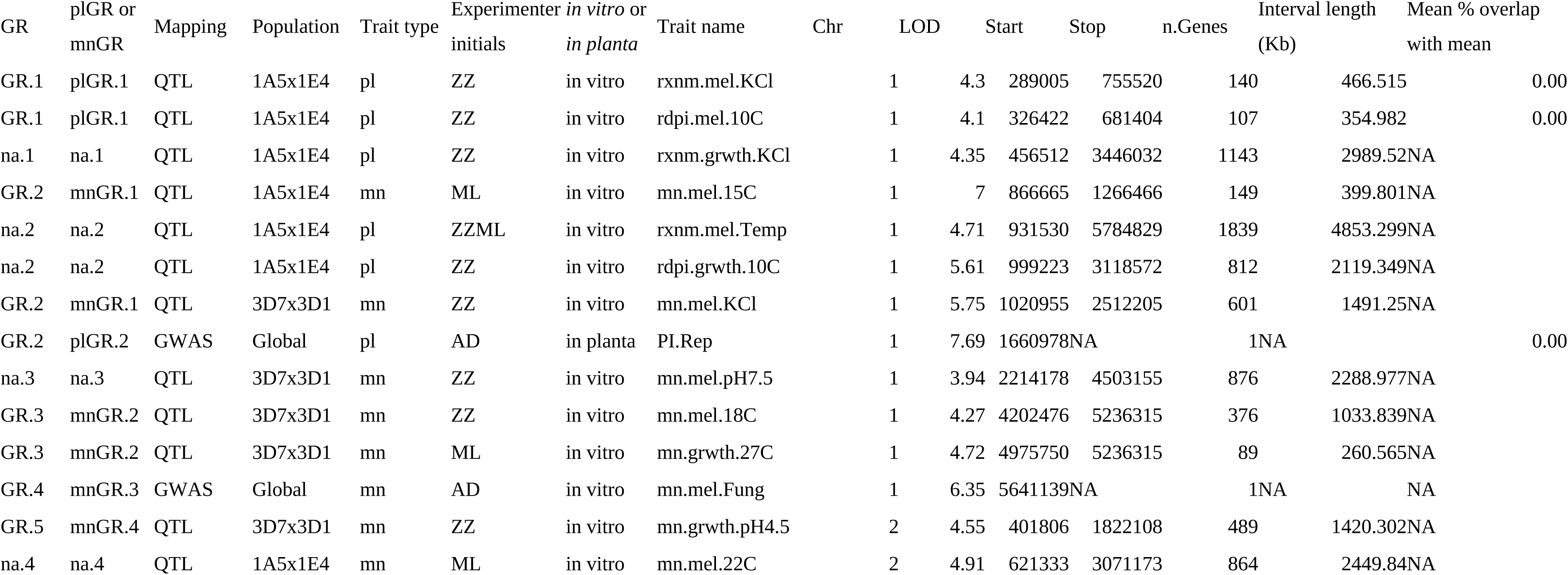
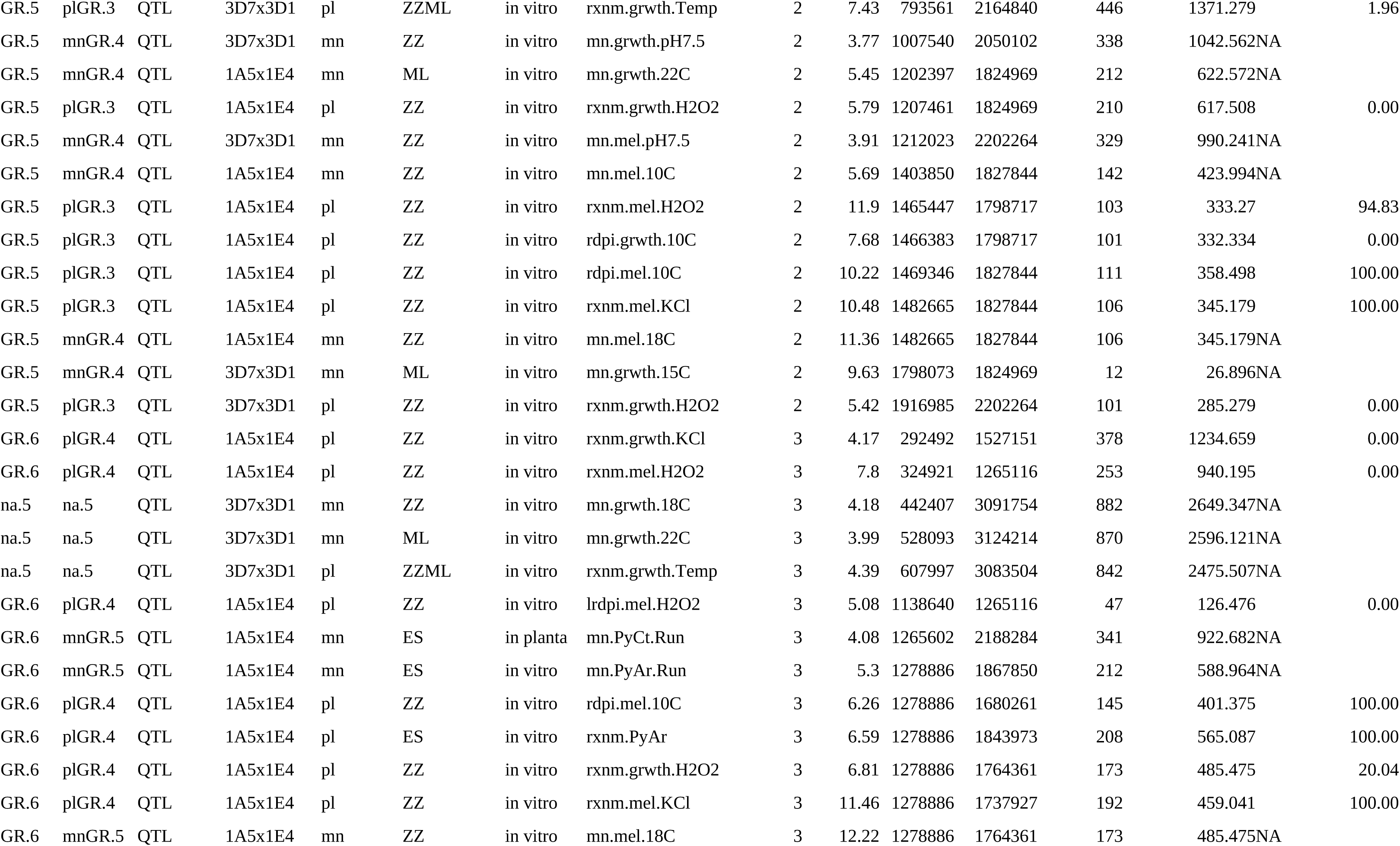
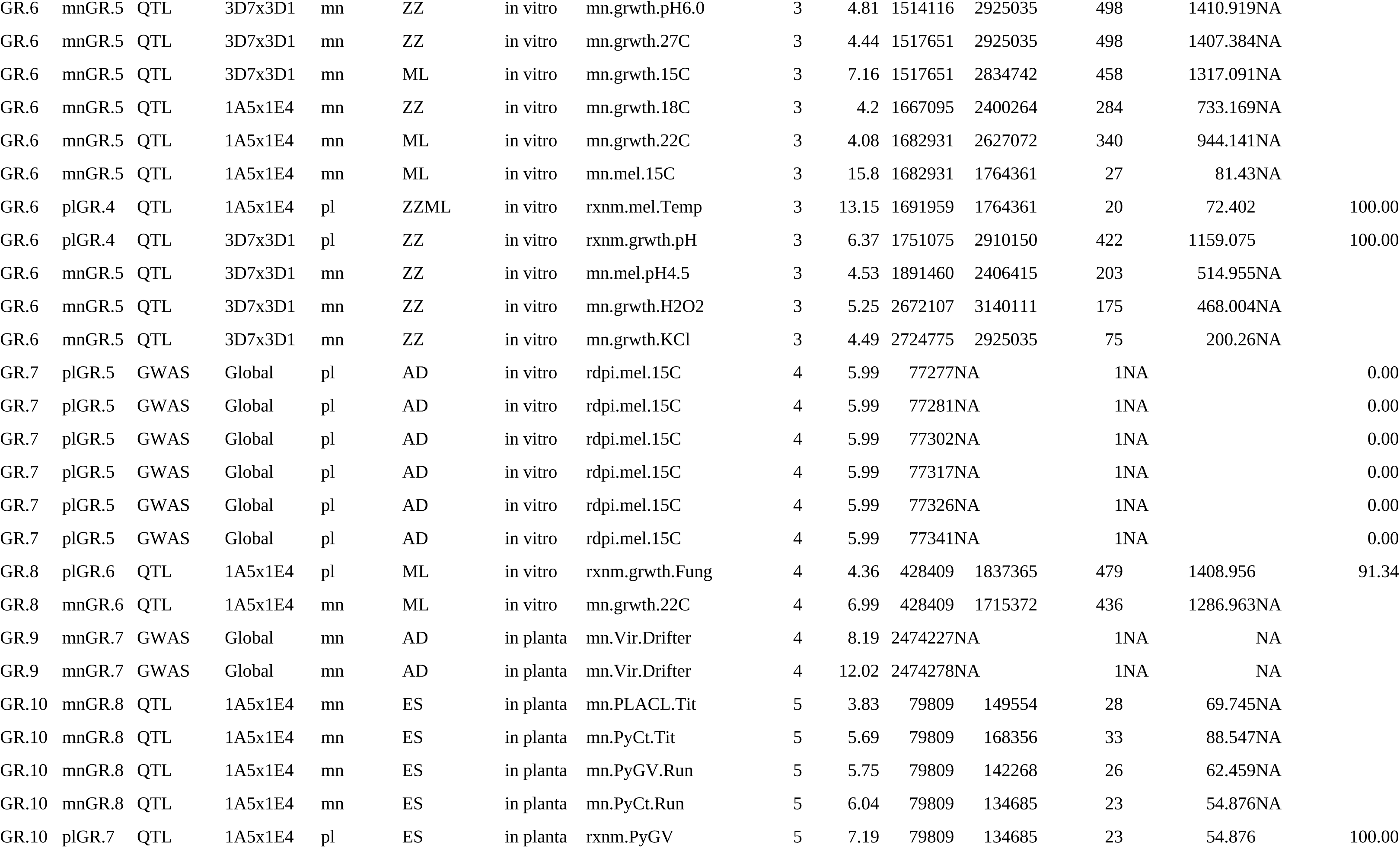
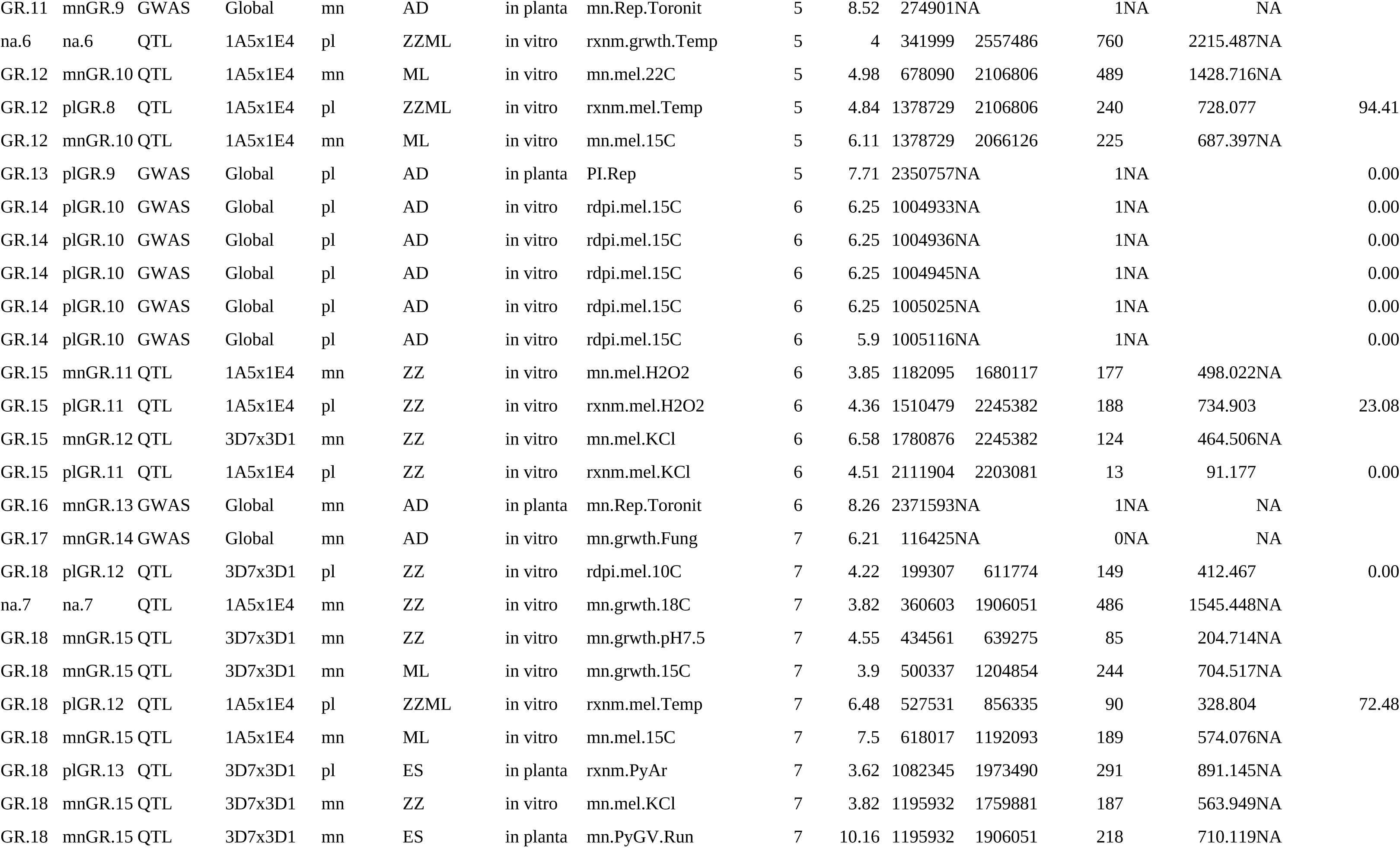
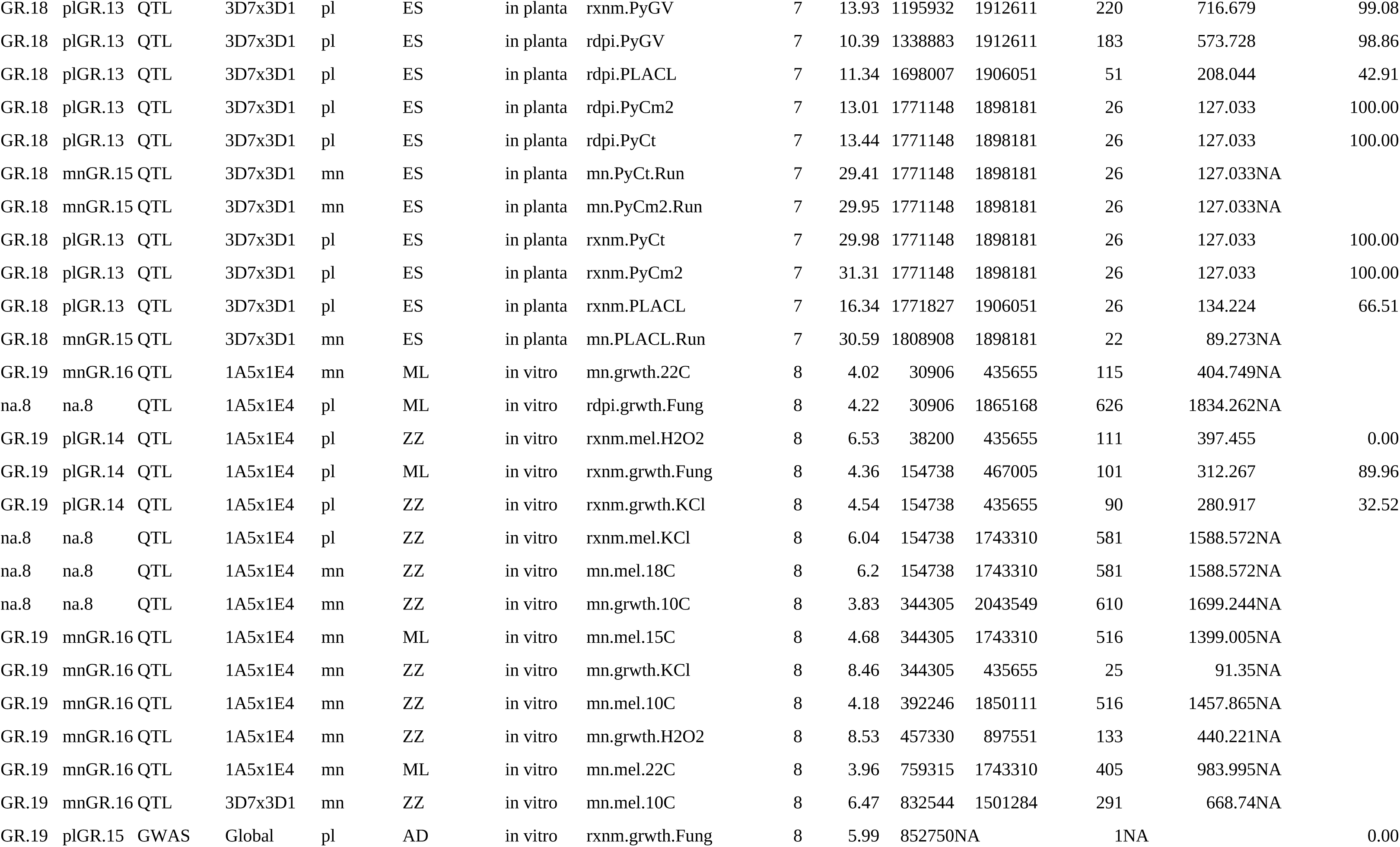
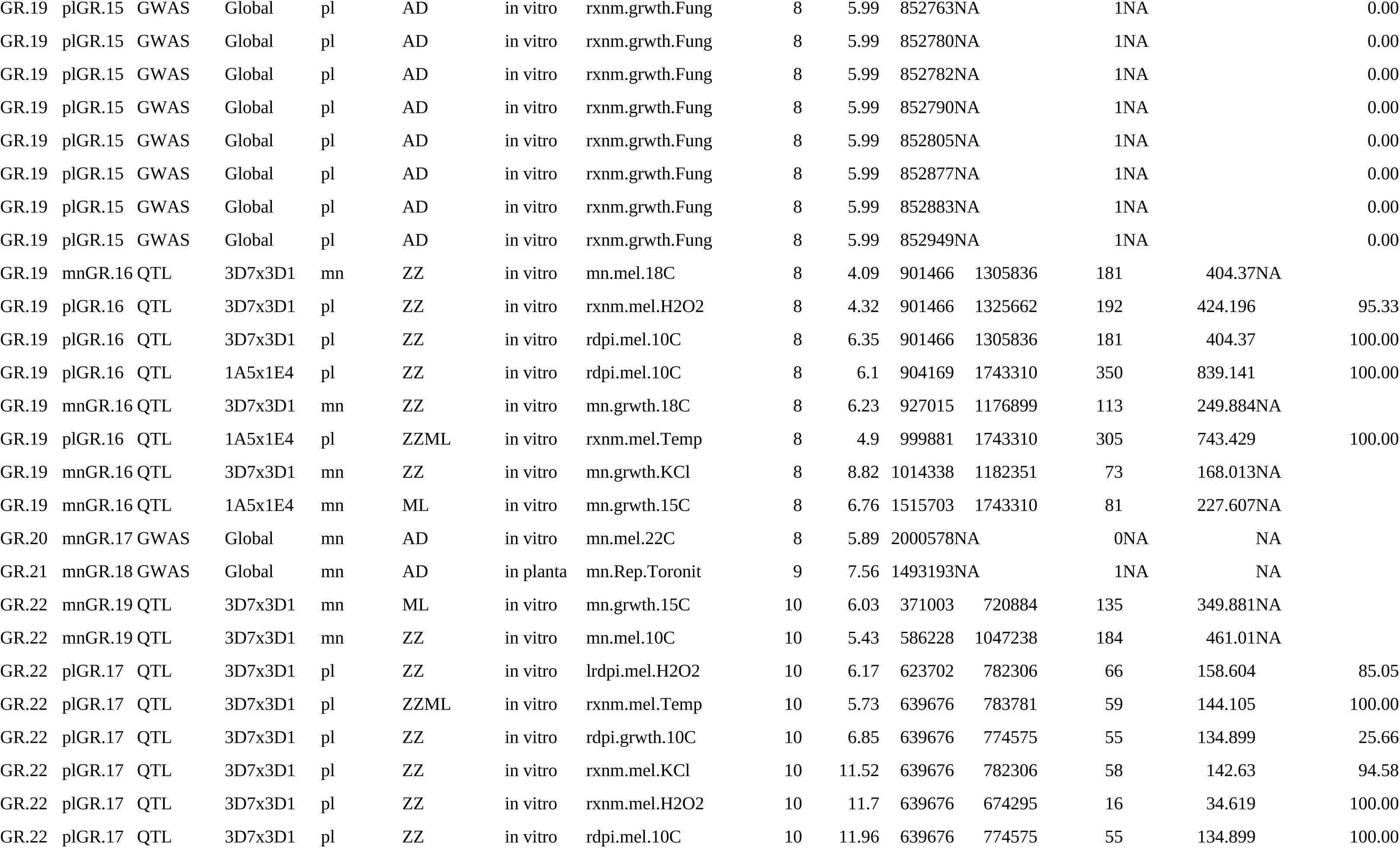
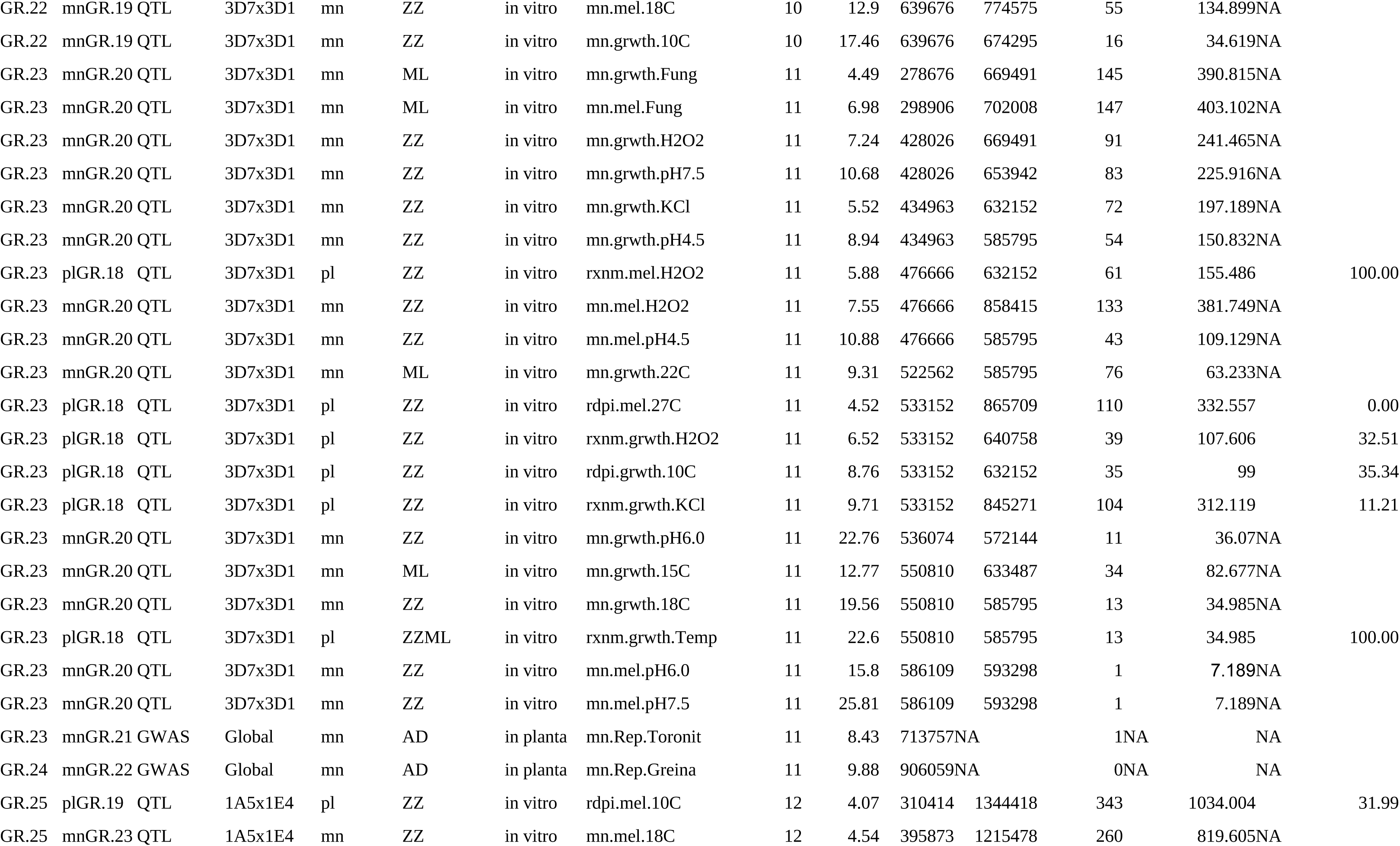
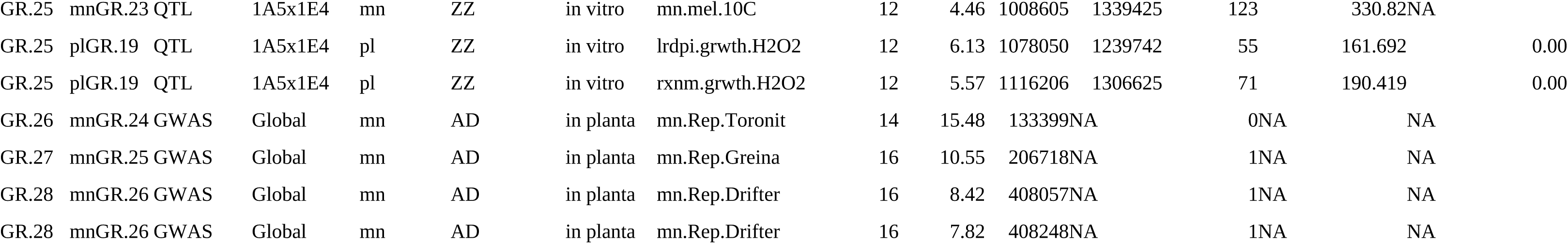
A complete list of significant QTL intervals and GWAS SNPs identified in these analyses. Each significant association includes the mapping method (QTL,GWAS), the experimental population used for the analysis, the experimenter who collected the data, the type of trait (pl: plasticity, mn:mean), an abbreviated trait name, the chromosome location, the QTL LOD score or GWAS -log(p-value) associated with the trait, the genomic position in the IPO323 reference genome, the number of IPO323 genes in the interval and the combined length of the genes, and the mean percentage of overlap between a plasticity QTL and the corresponding mean QTL. Overlapping QTL across all traits were collapsed into genomic regions (GRs), with some GRs specific for plasticity (plGR) and other GRs specific for trait means (mnGR). Large QTL intervals (>1500Kb) were excluded when defining the genomic regions (indicated with na.1, na.2, etc.).

To investigate the overlap between all QTL and GWAS SNPs, we removed 14 QTL intervals that were larger than 1500 Kb (ranging from 1545.4-4853.3 Kb in size) to reduce the number of false positive overlaps. After removing these large intervals there were 28 distinct genomic regions (GRs) harbouring either overlapping QTL (11 GRs) or GWAS SNPs (14 GRs) or both (3 GRs) (Supplementary Table S4a). The genomic regions containing only GWAS SNP(s) carried between 1-9 SNPs (covering 0-199 base pairs) and the QTL interval lengths ranged from 34.6 - 1409 Kb.

Considering only the plasticity traits that remained after removing the large intervals; among 57 plasticity traits tested we identified 56 QTL (29 in the 1A5x1E4 population; 27 in the 3D7x3D1 population) for 22 different traits, and 22 significant GWAS SNP associations for three different traits (one *in planta*, two *in vitro*). After merging overlapping intervals, we found 19 distinct plasticity genomic regions (plGR) (Supplementary Table S4b). We did not find QTL for any of the plasticity index (PI) traits – the plasticity measure across multiple environments (range 2-11), but we found significant GWAS SNPs (GR.2, GR.13) for plasticity index for reproduction measured across eight wheat cultivars (*in planta)*. These results suggest that within our QTL populations there were no large effect loci controlling plasticity across multiple environments, but there were a few SNPs associated with PI across eight host environments.

Among the 54 trait mean values (QTL:32, GWAS:22) we identified 71 QTL (28 in the 1A5x1E4 population, 43 in the 3D7x3D1 population) for 28 different traits, and 14 GWAS SNPs for seven traits (three *in vitro*, four *in planta*). After merging overlapping intervals, we found 26 distinct trait mean genomic regions (mnGR) (Supplementary Table S4c).

Many QTL intervals overlapped with other QTL intervals, and three GWAS SNPs were found within QTL intervals (GR.2, GR.19, GR.23; Figure 1). Next we describe these findings in the context of the five questions described in the methods.

### Q1) Do plasticity QTL or GWAS loci overlap with QTL or GWAS loci for the mean of the corresponding trait?

There was considerable similarity between the genetic architecture of trait plasticity and the genetic architecture of the mean trait value in the QTL analyses (Figure 1, Table 3, full details Supplementary Table S5), however no overlaps were found in the GWAS analyses. We propose two possible explanations for this pattern: 1) the GWAS associations often included only one or a few SNPs, so the intervals were very narrow, while the QTL analyses identified chromosome intervals that were much larger, increasing the likelihood of some overlap; 2) the GWAS included only a few plasticity traits and a modest number of strains, which limited the power to detect significant associations; only three plasticity traits were mapped in the GWAS. Considerable overlap between plQTL or plGWAS associations with the mnQTL or mnGWAS associations is expected given the correlations between phenotypic traits – the ACC is generally higher for a plasticity trait with its corresponding mean compared to all pair-wise correlations, except for the global *in planta* dataset (Figure S2). Consistent with this, we found that the overlaps between plQTL and the corresponding mnQTL trait were generally more highly skewed (75-100%), compared to other subsets of QTL intervals or all pairwise comparisons (Supplementary Figure S5).

**Table 3.** Percentage of plQTL or plGWAS-SNP (collectively called pl.Loc) overlapping with one of the corresponding mnQTL or mnGWAS-SNP (collectively called mn.Loc) for each plasticity trait. Each plasticity trait has between 2-8 corresponding mean traits. Each trait can have more than one significant QTL or GWAS-SNP association.

| Population | Plasticity trait | Number of<br>pl.Loc | Number of<br>mn.Loc | Number of<br>pl.Loc with<br>overlapping<br>mn.Loc | Percentage of<br>pl.Loc with<br>overlapping<br>mn.Loc |
| --- | --- | --- | --- | --- | --- |
| 1A5x1E4 | lrdpi.grwth.H2O2 | 1 | 2 | 0 | 0 |
| 1A5x1E4 | lrdpi.mel.H2O2 | 1 | 5 | 0 | 0 |
| 1A5x1E4 | rdpi.grwth.10C | 1 | 3 | 0 | 0 |
| 1A5x1E4 | rxnm.grwth.H2O2 | 3 | 7 | 1 | 33.33 |
| 1A5x1E4 | rxnm.grwth.KCl | 2 | 4 | 1 | 50 |
| 1A5x1E4 | rxnm.mel.H2O2 | 4 | 20 | 2 | 50 |
| 1A5x1E4 | rxnm.mel.KCl | 4 | 16 | 2 | 50 |
| 1A5x1E4 | rdpi.mel.10C | 5 | 35 | 4 | 80 |
| 1A5x1E4 | rxnm.PyAr | 1 | 1 | 1 | 100 |
| 1A5x1E4 | rxnm.PyGV | 1 | 1 | 1 | 100 |
| 1A5x1E4 | rxnm.grwth.Fung | 2 | 2 | 2 | 100 |
| 1A5x1E4 | rxnm.mel.Temp | 4 | 60 | 4 | 100 |
| 3D7x3D1 | rxnm.PyAr | 1 | 0 | NA | NA |
| 3D7x3D1 | rdpi.mel.27C | 1 | 3 | 0 | 0 |
| 3D7x3D1 | rxnm.grwth.H2O2 | 2 | 9 | 1 | 50 |
| 3D7x3D1 | rdpi.mel.10C | 3 | 15 | 2 | 66.67 |
| 3D7x3D1 | lrdpi.mel.H2O2 | 1 | 4 | 1 | 100 |
| 3D7x3D1 | rdpi.PLACL | 1 | 1 | 1 | 100 |
| 3D7x3D1 | rdpi.PyCm2 | 1 | 1 | 1 | 100 |
| 3D7x3D1 | rdpi.PyCt | 1 | 1 | 1 | 100 |
| 3D7x3D1 | rdpi.PyGV | 1 | 1 | 1 | 100 |
| 3D7x3D1 | rdpi.grwth.10C | 2 | 8 | 2 | 100 |
| 3D7x3D1 | rxnm.PLACL | 1 | 1 | 1 | 100 |
| 3D7x3D1 | rxnm.PyCm2 | 1 | 1 | 1 | 100 |
| 3D7x3D1 | rxnm.PyCt | 1 | 1 | 1 | 100 |
| 3D7x3D1 | rxnm.PyGV | 1 | 1 | 1 | 100 |
| 3D7x3D1 | rxnm.grwth.KCl | 1 | 6 | 1 | 100 |
| 3D7x3D1 | rxnm.grwth.Temp | 2 | 26 | 2 | 100 |
| 3D7x3D1 | rxnm.grwth.pH | 1 | 7 | 1 | 100 |
| 3D7x3D1 | rxnm.mel.H2O2 | 3 | 12 | 3 | 100 |
| 3D7x3D1 | rxnm.mel.KCl | 1 | 6 | 1 | 100 |
| 3D7x3D1 | rxnm.mel.Temp | 1 | 5 | 1 | 100 |
| Global | PI.Rep | 2 | 18 | 0 | 0 |
| Global | rdpi.mel.15C | 1 | 1 | 0 | 0 |
| Global | rxnm.grwth.Fung | 1 | 1 | 0 | 0 |

In total 56 plasticity QTL were mapped and for all but one of these we also mapped at least one of the corresponding mnQTL. The exception was in the 3D7x3D1 population, where a QTL for the plasticity trait rxnm.PyAr was found but no QTL were found for either corresponding mean (mn.PyAr.Run or mn.PyAr.Tit). There was no difference between the LOD or the interval length between a plasticity QTL and the corresponding mean QTL (LOD: Wilcox paired test V =595.5, *p*=0.20, interval length: Wilcox paired test V =410.5, *p*=0.15). In total, plQTL overlapped with at least one of the corresponding mnQTL 75% of the time (41 out of 55; 18 of 29 (62%) in the 1A5x1E4 population; 23 of 26 (88%) in the 3D7x3D1 population) (for all details see Supplementary Table S5). Fourteen plQTL did not overlap with at least one of the corresponding mnQTL, even though at least one mnQTL was mapped in each case. This suggests that 25% (14/55) of our plQTL map to different genomic regions than the mnQTL. The mnACC between plasticity traits and mean traits that have overlapping QTL was not different for plQTL that do not overlap the mnQTL (Supplementary Table S5a and S5b; mnACC overlapping =0.65, mnACC non-overlapping =0.52; Wilcox test: W = 351.5, p-value = 0.07). Summarising overlap by the number of traits is less straightforward because traits can map to multiple genomic regions (plQTL), however we can estimate the percentage of plQTL that overlap the mnQTL for each trait (Table 3). Considering plasticity traits with more than one plQTL (12 traits), in five cases all plQTL for that trait overlapped a mnQTL (100% of plQTL overlapped mnQTL); for example RXNM for melanisation rate across variation in temperature had four plQTL and all four overlapped at least one mnQTL. For the other seven traits with multiple QTL, overlap ranged from 30-80%. At the other end of the spectrum we identified four traits in which only a single plQTL was found, and for these plQTL there were no overlaps with the mnQTL (1A5x1E4: lrdpi.grwth.H2O2 (Chr12, GR.25), lrdpi.mel.H2O2 (Chr3, GR.6), rdpi.grwth.10C (including a large interval excluded from analysis on Chr1 and Chr2 GR.5) and 3D7x3D1: rdpi.mel.27C (GR.23 Chr11)). The ACC between these four plasticity traits and their mean trait was significantly lower (mnACC=0.36) than the mnACC between traits with overlapping QTL (Wilcox test: W = 55, p-value = 0.002).

For the GWAS analysis we found 36 significant associations for three different plasticity traits. No overlap was found between the plGWAS-SNP and corresponding mnGWAS-SNP, even though we found significant GWAS-SNP associations for a plasticity trait and the corresponding mean trait (Table 3, Supplementary Table S5c). For example, RXNM in growth rate across variation in fungicide concentration (Chr8) mapped to a different genomic interval than the mean growth rate in the presence of fungicide on Chr7. RDPI in melanisation rate across variation in temperature (15-22^0^C) mapped to Chr4 and Chr6, whereas mean melanisation rate at 22^0^C mapped to Chr8. The PI in reproduction across eight hosts mapped to Chr1 and Chr5 and mean reproduction in Drifter and Toronit and Greina mapped to different genomic regions (Table 2).

### Q2) Is there overlap in QTL and GWAS SNPs?

Three GWAS SNPs were within QTL intervals when both mean and plasticity traits were considered. However, if we consider only plasticity traits, none of the GWAS SNPs associated with plasticity were located within a plQTL (Supplementary Table S4b). Considering all genomic regions, GR.2 (Chr1) contains two mean trait QTL from different experiments and populations (ML: 1A5x1E4, mean melanisation rate at 15^0^C, ZZ: 3D7x3D1 mean melanisation rate in the presence of KCl) and one GWAS SNP associated with PI in reproduction measured across eight hosts. In GR.19, there were nine GWAS SNPs significantly associated with the *in vitro* trait RXNM plasticity for growth rate across variation in fungicide concentration (rxnm.grwth.fung) mapping to Chr8:852750-852949. The same trait measured in the 1A5x1E4 population mapped to an interval 417095 bp away (Chr8: 154738-467005). The GWAS SNPs fall within the QTL intervals for several mean traits ((mn.mel.15C, mn.mel.10C (1A5), mn.grwth.H2O2, mn.mel.22C, mn.mel.10C (3D7)), but do not fall within any plQTL. In GR.23 a GWAS SNP associated with mean reproduction on the host Toronit (mn.rep.Toronit) falls within the QTL interval of two *in vitro* mean traits (e.g. mn.mel.H2O2, rxnm.grwth.KCl).

### Q3) Is there overlap between QTL intervals across *in vitro* and *in planta* experiments?

There were five genomic regions (GR.2, GR.6, GR.18, GR.19 and GR.23) where QTL for traits measured *in vitro* overlapped with traits measured *in planta*. Two (GR.2, GR.19) were highlighted above. In GR.6 (Chr3), three *in planta* traits map to a region containing 18 QTL for *in vitro* traits. In GR.18 (Chr7), 12 pycnidia plasticity and mean traits measured across two cultivars map to the same genomic region as QTL for six *in vitro* traits (Figure 2). If we consider only plasticity traits, there was one genomic region (plGR.4) where associations for one *in planta* and eight *in vitro* traits overlap. These include *in vitro* RXNM and RDPI in melanisation and growth rate across variation in temperature, pH, salt concentration and hydrogen peroxide concentration overlapping with a QTL for RXNM for pycnidia area measured across two host genotypes. If we consider the phenotypic correlations between *in planta* and *in vitro* traits, these were not clustering, suggesting modest phenotypic correlations.

**Figure 2.**
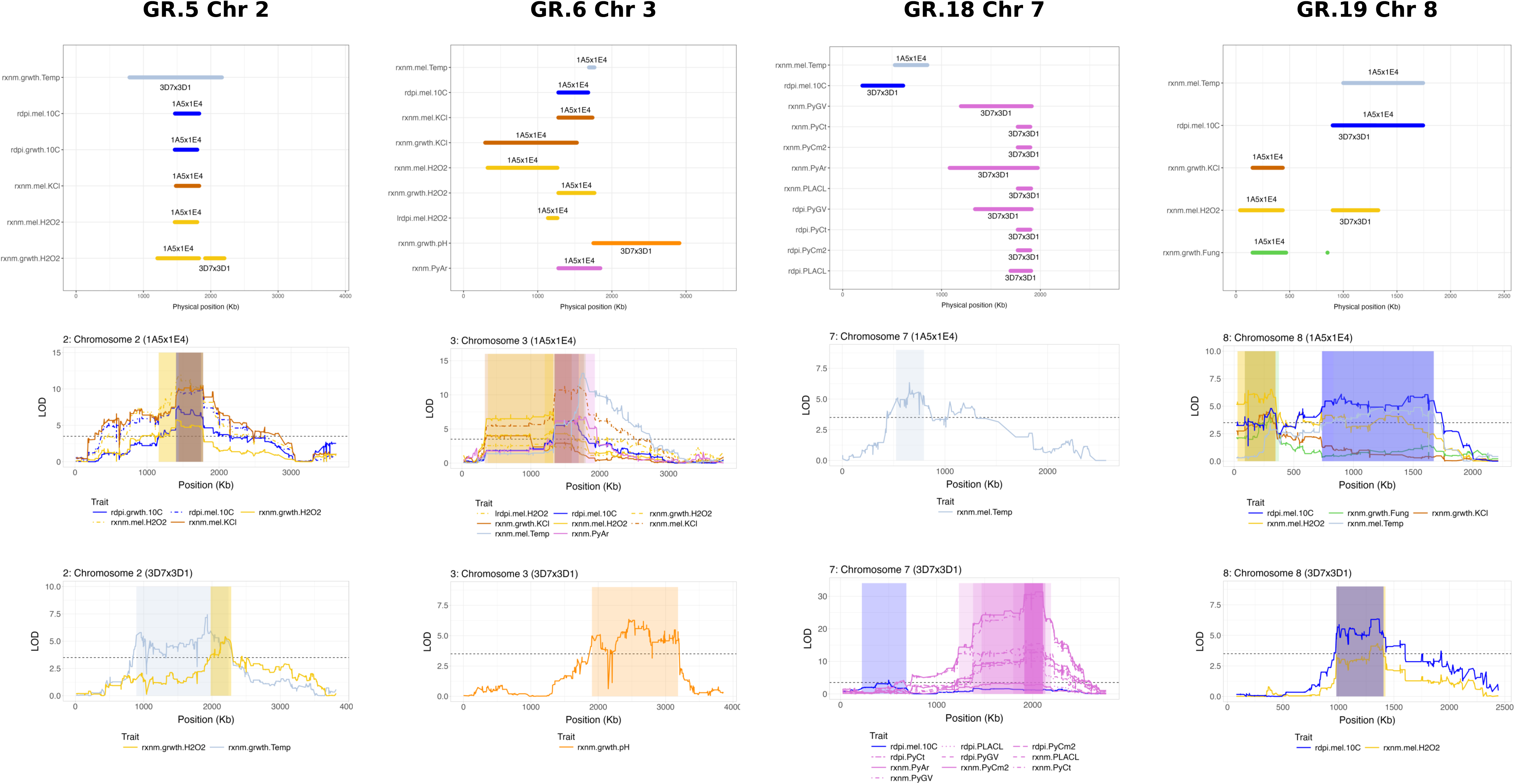
Genomic regions containing significant QTL and/or GWAS associations with plasticity traits. The top row of panels shows the plQTL intervals and plGWAS SNP genomic position (bp) on four chromosomes based on the IPO323 reference genome with the QTL population indicated above/below the line. The plGWAS SNP is not labeled. The middle row shows LOD plots for plQTL in the 1A5x1E4 cross with genomic position (bp) based on the ZT99_1A5 reference genome. The bottom row shows LOD plots for plQTL in the 3D7x3D1 cross with genomic position (bp) based on the ZT99_3D7 reference genome.

**Figure 3.**
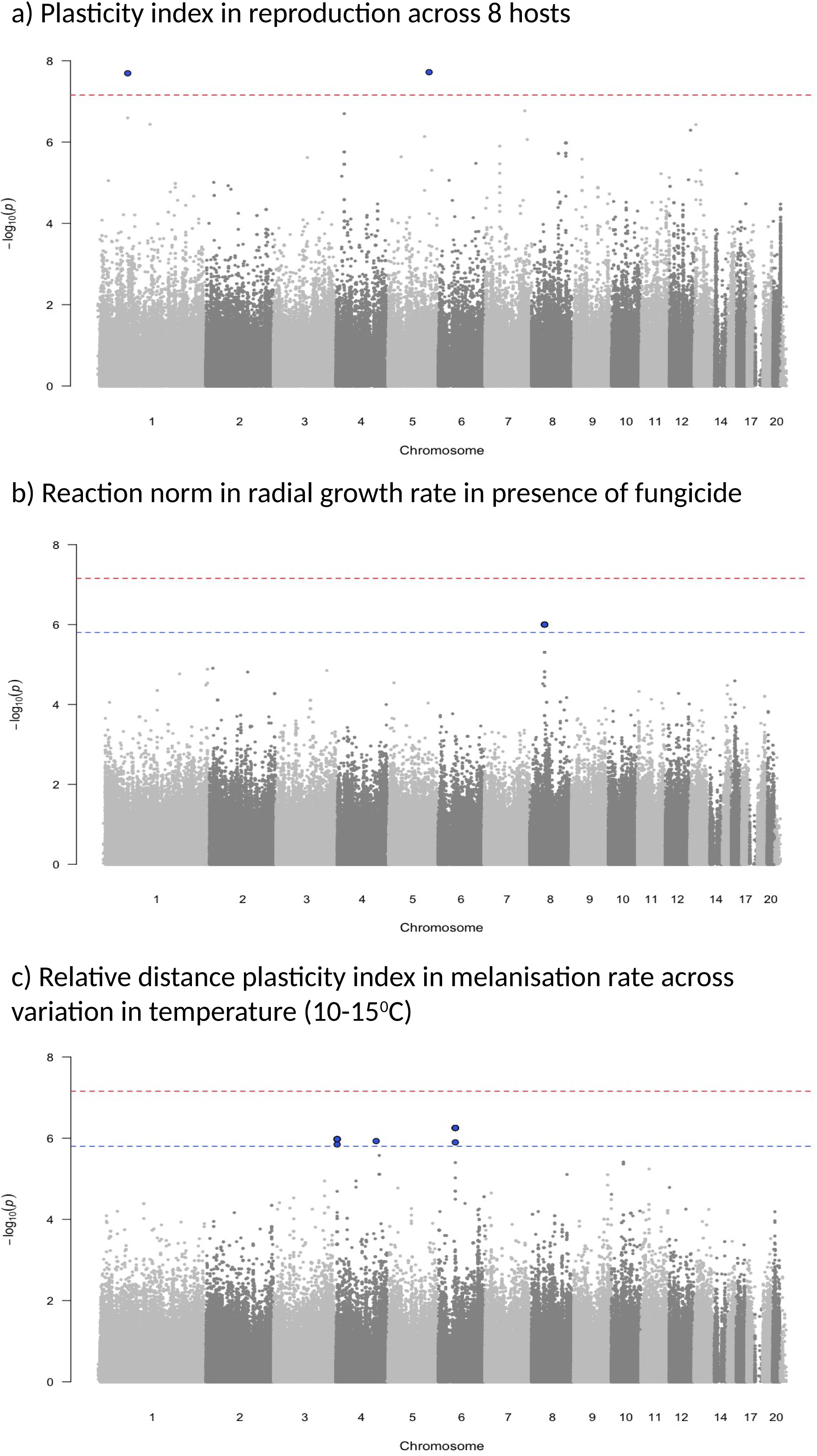
Manhattan plots of three plasticity traits showing significant GWAS associations for one *in planta* trait and two *in vitro* traits. Blue dashed lines are based on chromosome-level significance thresholds and red dashed lines are based on genome-wide significance thresholds. Significant SNPs are colored blue and enlarged.

### Q4) Is there overlap in QTL identified in the two experimental cross populations (1A5x1E4 and 3D7x3D1)?

Considering all traits, we identified six GRs (GR.2, GR.5, GR.6, GR.15, GR.18, GR.19) where traits measured in both populations overlap. Considering only plasticity, there were four plGRs (plGR.3, plGR.4, plGR.12, plGR.16). In GR.19 (Chr8), the same plasticity trait in both populations - RDPI in melanisation rate across 10-18^0^C, maps to essentially the same genomic position (Table 2, Figure 2). In GR.5 the same trait (RXNM for growth rate across variation in reactive oxygen) maps to adjacent QTLs separated by only 92016 bp (1A5x1E4 Chr2:1207461-1824969, 3D7x3D1 Chr2: 1916985-2202264).

### Q5) Is there overlap among QTL for plasticity measured across environments?

Of the 13 plGRs with more than one plQTL, nine contained QTL for plasticity traits measured across variation in abiotic (KCl, H_2_O_2_, pH, temperature, fungicide) or biotic (two hosts) environments. In four of these plGRs (plGR.3, plGR.4, plGR.17, plGR.18), plQTL associated with variation across at least three different environments overlapped (Supplementary Table S4b). In plGR.4 (Chr3), plasticity in growth/melanisation across variation in salt, reactive oxygen concentration, temperature and pH, and pycnidia area across two hosts all mapped to a single genomic region. All but one of the plasticity traits were measured in the 1A5x1E4 population. This matches somewhat with the pattern of clustering of traits based on the phenotypic correlations within 1A5x1E4, which found clustering among melanisation and growth across variation in salt, reactive oxygen and temperature, however plasticity across variation in fungicide concentration and pycnidia area were not strongly correlated with these other plasticity traits (Supplementary Figure S2).

### Identifying candidate genes for plasticity in selected regions of interest

We present next the results of three approaches used to identify candidate genes for trait plasticity: (a) GO enrichment analysis of identified genomic regions; (b) Biosynthetic gene cluster enrichment and; (c) a focus on narrow chromosome intervals with high LOD values.

a. **Go enrichment approach**

We performed a GO enrichment analysis using the genes from all significant QTL intervals and genes with significant GWAS SNPs. If we exclude the large QTL intervals (>1.5 MB), we have a list of 6675 genes spread across 13 chromosomes. From this analysis we identified one enriched GO term for Biological Process (BP), namely GO:0043386: mycotoxin biosynthetic process. No enriched GO terms for Cellular Component (CC) or Molecular Function (MF) were significant (Supplementary Table S6a). Considering intervals containing QTL for traits measured across more than two environments (seven GRs: GR.2, 5, 6, 17, 19, 22, 23) there were 3585 genes. No enriched GO terms were identified (Supplementary Table S6b) in these regions.

If we restrict the analysis to genes within plasticity QTL (plGR), there were 4353 genes spread across 11 chromosomes. From this analysis we identified one enriched GO term for BP, namely GO:0043386: mycotoxin biosynthetic process, and one enriched GO term for CC - GO:0008236: serine-type peptidase activity. No MF GO terms were significant (Supplementary Table S6c). Considering intervals containing plQTL for traits measured across more than two environments (five plGRs: plGR.3, 4, 14, 17,18) there were 1835 genes. We identified no enriched GO terms for BP or CC, but for MF there were three: GO:0003887: DNA-directed DNA polymerase activity; GO:0036402: proteasome-activating activity, GO:0008236: serine-type peptidase activity (Supplementary Table S6d).

**Biosynthetic gene cluster overlap**

Of the 28 GRs, 11 contained >1 BGC (Supplementary Table S4a). The GRs with the most BGCs were GR.5 on Chr2 and GR.23 on Chr11, which each contained three BGCs. In the following enrichment analysis we excluded GRs containing only GWAS SNPs. The number of BGCs found within observed GRs was greater than the number found within simulated intervals (the median number of BGC (mdBGC) in real GR = 17, the number found in simulated GR mdBGC = 11, 95%CI (6, 17), p=0.028). If we consider each individual GR separately, we found only one GR enriched for BGC, GR.23 Chr11, which contains three BGCs (p=0.03). This GR contained significant associations with traits measured across five different environments (19 QTL and one GWAS SNP) (Supplementary Table S4a). Considering plGRs, 6 from 19 plGR contained >1 BGC (Supplementary Table S4b). The number of BGCs found in real plGR was not different from the number found in simulated plGR (the mdBGC in real plGR = 9, in simulated plGR mdBGC = 9, 95%CI (4, 15), p=0.61). No single plGR was significantly enriched for BGCs, however plGR.18 on Chr 11, which had two BGCs, approached significance (p=0.06, median=0, 95%CI (0,2)). Considering the GRs and plGRs containing only GWAS SNP(s), an intergenic GWAS SNP in GR.24 on chromosome 11 occurred within a NRPS-Like BGC. The other GWAS SNP found within a BGC was on GR.21 on chromosome 9, with the SNP within a gene encoding a DNA-directed RNA polymerase (ZtIPO323_101960) and the next gene encoding a polyketide synthase (*Pks*) (ZtIPO323_101970).

**Narrow intervals with high LOD values**

The highest LOD recorded (31.3) was for a QTL located on Chr7 (GR.18, plGR.13). This GR contained five QTL with high LOD (29.4-31.3, Table 2). Plasticity across two host genotypes (Titlis and Runal) in pycnidia traits and virulence (PLACL) mapped to this region in the 3D7x3D1 population. The specific QTL with the highest LOD was for RXNM in pycnidia density (rxnm.PyCm2) and contained 26 predicted genes. This region contains *Avr3D1*, which encodes a small secreted protein that triggers quantitative resistance in cultivars with the *Stb7* resistance gene (Meile et al., 2018) (Supplementary Table S7).

The genomic region containing a plQTL with the next highest LOD (22.6 for rxnm.grwth.Temp) was GR.23 (plGR.18) on Chr11 in the 3D7x3D1 population (Table 2). The GR.23 interval is enriched for BGCs, including two non-ribosomal peptide synthetase (NRPS)-Like clusters (with 12 and 30 genes respectively) and one *Pks* cluster (with 14 genes). A total of six plasticity traits measured across different environments mapped to this genomic region. RXNM in growth rate across variation in temperature (10-27^0^C) had the highest LOD and contained only 13 genes (Supplementary Table S7), between ZtIPO323_112260-ZtIPO323_112410. The interval contains a putative 26S proteasome regulatory subunit rpn9 gene (ZtIPO323_112330), that was shown to play a role in temperature sensitive growth in *S. cerevisiae* (Takeuchi et al. 1999). This gene is also associated with the enriched GO term proteasome-activating activity. This interval also contains *Mog1* (ZtIPO323_112320). Variation in *Mog1* also affects temperature sensitive growth in *S. cerevisiae* (Baker et al. 2001) and is also involved in response to osmotic stress (Kelley and Paschal 2007). The 95% confidence interval for this QTL ends within the immediately following *Pks1* gene (ZtIPO323_112420). *Pks1* encodes a polyketide synthase involved in DHN melanin biosynthesis (Jia et al., 2021). A high LOD QTL interval for mean traits (mnQTL.20) also maps to this region. The QTL interval for mean melanisation rate at pH7.5 and pH6.0 (LOD =25.81 and 15.8 respectively) fall within this *Pks1* gene. This interval was only 49 bp wide in the ZT993D7 genome, and mean growth rate traits (pH6.0, 18^0^C) mapped to essentially the same region as rxnm.grwth.temp, with high LOD (22.76, 19.56 respectively). In a previous QTL analysis Lendenmann et al (2014) proposed that two nonsynonymous SNPs (affecting amino acid positions 884 and 1783) within *Pks1* could be responsible for variation in melanisation rate. The peak marker within the very small (49 bp) interval we identified is the nonsynonymous SNP that alters amino acid 884 from a valine into alanine (ZT3D7 Chr11:658119).

Multiple QTL from 1A5 mapped to an overlapping region (GR.5, plGR.3) on Chr2 with LOD>10 (Table 1, Figure 2). This region contained three BGCs, including two NRPS BGCs (with 14 and 13 genes respectively) and one terpene BGC (31 genes). Considering only the higher (>10) LOD QTLs, there was RXNM in melanisation rate across variation in H_2_O_2_ and salt concentration, and RDPI in melanisation rate across temperature variation (10-18^0^C) in the 1A5x1E4 population. The highest LOD (11.9) QTL in the 1A5x1E4 population for RXNM plasticity in melanisation rate across variation in H_2_O_2_ has 123 genes. This QTL does not overlap with a BGC, but it contains several interesting candidate genes that we list in genomic order. The first encodes a heat shock protein, *Hsp98* (ZtIPO323_029190), the second is a gene encoding an UV radiation resistance protein (ZtIPO323_029520), the third gene encodes an oxidoreductase like protein (ZtIPO323_029800) and the fourth includes two genes encoding prefoldin subunit4, which is related to HSPs (Supplementary Table S7).

Another notable region with overlapping QTLs and LOD>10 (GR.6, plGR.4) was found on Chr3 (Table 1). Eleven plasticity traits (including the large intervals that were excluded when calculating overlap) measured *in vitro* and one measured *in planta* (RXNM pycnidia area) mapped to this interval. The QTL intervals from the 3D7x3D1 population were relatively large (422-842 genes) compared to the intervals in the 1A5x1E4 population (>266 genes). Two of the QTL in the 1A5x1E4 population, RXNM in melanisation rate across variation in temperature and salt concentration, have a LOD>10. The interval with the highest LOD (13.15, rxnm.mel.temp) has only 20 genes. Of the 14 genes in this interval with predicted functional domains, eight (57%) have putative functions in fungal stress responses. The most notable is the gene encoding the Woronin body major protein, *Hex-1*. This gene encodes known pleiotropic responses involving vegetative growth, osmotic and reactive oxygen stress responses, and pathogenicity in plant pathogens (Vangalis et al. 2020). The interval also contains a gene encoding a glycoside hydrolase family 17 protein (a CAZyme), which is a secreted protein that can act as a virulence factor (Bradley et al. 2022). If we consider the larger interval, with the second highest LOD =11.46 (rxnm.mel.KCl), a possible candidate is a gene encoding a heat shock factor binding protein 1 (Hsf1). This protein is considered a master regulator of heat shock response, activating the heat shock proteins under proteotoxic stress caused by heat and oxidative stress (Supplementary Table S7).

In the 3D7x3D1 population, multiple measures of plasticity in melanisation rate in response to variation in H_2_O_2_ and salt concentration and variation in temperature (10-18^0^C) all map to an overlapping region (GR.22, plGR.17) on chromosome 10 (Table 1). This GR contains two BGCs, including one terpene and one PKS-NRPS, containing 29 and 20 genes respectively. Three of the plQTL have LOD>10: plasticity in melanisation rate in response to variation in temperature (10-18^0^C) and reactive oxygen and salt concentration. The smallest interval (rxnm.mel.H2O2) has only 16 genes and does not contain a BGC. A mean trait QTL with high LOD (17.46, mn.grwth.10C) overlaps perfectly with this interval, and the interval overlaps with a previously identified QTL for cold sensitivity (Lendenmann et al., 2016), in which Pbs2, which phosphorylates the HOG1 pathway gene, was proposed as the candidate gene.

### Other noteworthy intervals containing known or plausible candidate genes

GR.19 on Chr8 contains QTL for plasticity traits from both populations as well as GWAS SNPs from the global population (Table 1). The same trait – RDPI in melanisation rate across temperature variation (10-18^0^C) - mapped to essentially the same interval (Figure 2, 1A5x1E4: Chr8:904169-1743310, 3D7x3D1: Chr8:901466-1305836) in both progeny populations, though the LOD scores were not especially high (6.1, 6.25). This interval has 195 genes including several plausible candidate genes. The first is a gene encoding a “Low temperature viability protein” (*Ltv1*), the second encodes a heat shock transcription factor and the third is *Ste20* – encoding a MAP kinase kinase that forms part of the HOG pathway. This GR.19 interval also contains nine GWAS SNPs (Chr8:85275-852949) for plasticity in growth rate across fungicide concentrations, and a QTL interval for the same trait. The GWAS SNP was in a gene encoding an amino acid permease protein (ZtIPO323_091180, Chr8:850630-853505) (Supplementary Table S8).

Another notable genomic region is GR.10 on Chr5. It contains a QTL for RXNM plasticity in pycnidia grey value across variation in two hosts, Titlis and Runal, and four mean pycnidia traits measured in the 1A5x1E4 population. This plQTL contains 23 genes, including the avirulence factor *AvrStb6* (ZtIPO323_060700). It also overlaps a Pks BGC, with the *Pks* gene and other key genes encoding enzymes and transcription factors.

Plasticity in growth rate across variation in H_2_O_2_ and plasticity in melanisation rate across variation in temperature (10-18^0^C) mapped to an interval on Chr12 (plGR.19). Within this interval is the gene *ZtMsr1,* a transcription factor controlling a temperature sensitive yeast-hypha morphological transition (Francisco et al. 2023). It also contains TFIID-18kDa-domain-containing protein, which coordinates transcription initiation.

## Discussion

This study genetically mapped multiple *in vitro* and *in planta* plasticity traits in a fungal plant pathogen using datasets that were generated across different projects over more than a decade of research. The study is notable because of the wide variety of environments that were included during phenotyping and because it quantifies the overlap between plasticity traits measured in different experimental populations, across different environmental gradients and between plasticity and the corresponding mean trait. This comprehensive approach provided unprecedented insights into the genetic architecture of trait plasticity in a plant pathogen.

We found that plasticity is a quantitative genetic trait that can be mapped to multiple regions in the genome. We also found that in most of the cases (75%) the QTL for plasticity in a trait overlapped with the QTL for the corresponding trait mean. This suggests that in most cases trait plasticity results from pleiotropy, the action of tightly linked loci, or both. There were four plasticity traits that mapped to separate genomic regions that did not overlap with the corresponding trait means, indicating that these traits are likely controlled by separate loci, and providing strong support for a role for epistasis in the plasticity of these traits. The GWAS analyses yielded fewer significant associations, possibly due to a limited power to detect associations resulting from the relatively small number of pathogen strains used in this dataset. In the GWAS analyses, no SNPs significantly associated with a plasticity trait overlapped with SNPs significantly associated with the corresponding mean. Another important finding is that plasticity measured across different environments can map to the same region of the genome. In one case plasticity measured across variation in five different environments (including *in vitro* and *in planta* traits) mapped to a single 2617 Kb genomic interval (plGR.4) on chromosome 3. This suggests that plasticity across multiple environments can be governed by the same gene (i.e. a master regulatory gene) or a genetically linked cluster of genes.

The QTL results are consistent with variation in plasticity resulting mainly from pleiotropy, whereas the GWAS results indicate a larger role for epistasis. Similar patterns have been observed in other studies. For example, QTL and GWAS studies in maize found partial overlaps between the mnQTL and plQTL (Jin et al., 2023; Tibbs-Cortes et al., 2024), whereas a different GWAS study in maize found that only four out of 977 SNPs were associated with both the trait mean and trait plasticity (Kusmec et al., 2017). These different outcomes may reflect differences in mapping precision: QTL intervals are typically much larger than GWAS intervals, although some of our QTL intervals were relatively narrow (one was only 7189 bp long and several had fewer than 50 genes, Table 1). The differences between the QTL and GWAS results may also reflect a lack of statistical power in our GWAS datasets. At least one variant must be strongly associated with two different phenotypes to observe co-localization and it is possible that the lack of co-localization observed in our GWAS analysis is due to a lack of statistical power. Our QTL studies had greater power to detect associations because we mapped more traits in a greater number of individuals (>250 offspring), and because the parents used in the QTL populations were chosen to be divergent in many of the measured traits (Lendenmann et al., 2014). In the QTL analysis we tested 72 traits and identified at least one QTL for 30 (41%) and 43 (59%) of the traits for the 1A5x1E4 and 3D7x3D1 populations, respectively. In contrast, for the global GWAS population, we mapped 16 *in vitro* traits and found no significant associations at a conservative genome-wide cut-off. When we used a less conservative chromosome-level cutoff we found at least one significant SNP association for five of these traits (31%). For the *in planta* datasets using the conservative genome-wide p-value cutoff, we found at least one significant SNP associated with five different traits (23%). Other factors such as trait heritability (h^2^) will also influence the probability of detecting significant associations, but h^2^ of traits used in the GWAS population were not lower than those measured in the QTL populations (Supplementary Figure S3). The phenotypic correlations between plasticity traits and the associated mean traits can reflect underlying genetic correlations – which are indicative of shared genetic control or pleiotropy. Across the GWAS *in planta* datasets, the ACC between a plasticity trait and the corresponding mean trait were generally lower (mACC =0.29) than the QTL traits, but the mACC between plasticity and mean traits for the *in vitro* GWAS dataset was similar to what was found for the QTL datasets (e.g. GWAS *in vitro*: mACC =0.37, QTL 1A5x1E4: mACC =0.37 and QTL 3D7x3D1: mACC =0.34). Without identifying the causal mutation through functional studies, we cannot determine with certainty if trait plasticity and the trait mean are controlled by the same gene (pleiotropy), or due to epistasis/interactions between tightly linked genes. From an evolutionary perspective, these two molecular mechanisms have the same outcome - selection on plasticity will also impact the trait mean and vice versa. In this case, there will be selective constraints because these traits cannot evolve independently.

The study also identified considerable overlap in plasticity measured across different environmental gradients and different populations. Multiple plQTL map to the same or overlapping genomic regions (plGR), suggesting a shared genetic control of plasticity across different environmental gradients and in different genetic backgrounds. Shared control of plasticity across environments could be mediated by highly pleiotropic genes, or master regulators, such as mitogen activated protein kinase (MAPK) pathways and heat shock factor proteins, or by hub genes within co-expression modules, for example biosynthetic gene clusters (BGC) (Baadani et al., 2025). Indeed, we found that genomic regions containing at least one QTL were enriched for BGC.

We identified several plausible candidate genes that could affect plasticity of the measured traits in our experimental populations. We first consider the plGR.4 interval on Chr3 that contains multiple plQTL measured across five different environmental gradients, including both *in vitro* and *in planta* traits. In the smallest, high LOD plQTL interval for rxnm.mel.temp, which contains 20 genes in a 72 kb interval, we found a gene belonging to the Woronin body major protein family containing a Hex-1 domain (Supplementary Table S7). The Woronin body is a specialized, membrane-bound peroxisome organelle found in filamentous ascomycete fungi. It is packed with a dense crystalline hexagonal peroxisomal protein, which supports the unique cellular architecture of the fungal hypha. Hex-1 is essential for sealing hyphal wounds to prevent leakage, but also has been shown to have pleiotropic effects on growth, asexual reproduction, stress response and virulence, although the effects can vary across different fungal pathogens (for a review see (Vangalis et al., 2020). For example, in *Aspergillus flavus* conidia formation and virulence were affected in *Hex-1* mutants, but the colony growth and stress response was similar to wildtype (Yuan et al., 2019). In *Ustilaginoidea virens*, UvHex-1 mutants could not infect rice, conidia production was slightly higher than the wildtype, and the mutants were more sensitive to salt and reactive oxygen stress (Mao et al., 2025). In *Verticillium dahliae*, deletion of *Hex-1* resulted in reduced conidia formation, slower growth in control conditions and under somatic and reactive oxygen stress and virulence was greatly reduced on eggplant seedlings (Vangalis et al., 2020). Deletion of *Hex-1* in *A. fumigatus* resulted in increased sensitivity to stressors that affect the cell wall integrity, for example 0.01% SDS, Calcofluor White, Congo Red and antifungal agents that target cell walls, however the *Hex-1* mutant response to osmotic stress was similar to the wildtype (Beck et al., 2013). There is Hex-1 coding sequence variation and differential expression among the parents used for QTL mapping. The coding sequence is identical between 1A5, 1E4, 3D7 and IPO323, but 3D1 carries a non-synonymous mutation (codon 128, Ser-Pro) (Supplementary Material SM10, (https://github.com/jessstapley/QTL-mapping-Z.-tritici/)). There was differential gene expression of the *Hex-1* 1A5 ortholog (ZT1A5_G3970) between 1A5 and 1E4; the gene was differentially expressed in colonies grown on minimal media *in invitro* and *in planta* at 28 dpi, but for the 3D7 ortholog (ZT3D7_3981) not enough reads mapped so differential expression could not be evaluated (https://github.com/jessstapley/QTL-mapping-Z.-tritici/). Based on the coding sequence analysis, it appears likely that the gene model in 3D7 is incorrect (Supplementary Material SM10). Although there is a perfect alignment in the Hex-1 portion of the gene, the RNA read mapping may have failed if the gene model was not correct in 3D7.Considering the pleiotropic nature of *Hex-1* in several fungal pathogens, we consider this to be a good candidate for regulating plasticity in growth, melanisation measured across variation in temperature, pH, salt and reactive oxygen concentration, and pycnidia area across host genotype. If we examine the second highest LOD plQTL in the plGR.4 interval (rxnm.mel.KCl), which is larger (192 genes), we find a well-known master regulatory gene (*Hsf*) encoding heat shock factor binding protein 1 (Hsf1). Hsf1 is a transcriptional regulator essential for a protective response to a diversity of stressors, including changes in temperature, pH, osmolarity and the presence of oxidizing agents (Veri et al., 2018), making it another plausible candidate gene to explain the observed plasticity. There is coding sequence variation across the QTL mapping parents and IPO323 in Hsf1. IPO323 and 3D7 have identical sequences, while 1A5, 1E4 and 3D1 have the same non synonymous mutation (codon22 Ser-Pro) (Supplementary Material SM11, (https://github.com/jessstapley/QTL-mapping-Z.-tritici/)). There were insufficient reads mapping to this gene in our RNAseq datasets to identify any differences in expression.

The plGR.4 interval is large (>2.6 Mb) and contains many genes, but it did not contain a BGC and it seems enriched for genes with roles in stress responses. 57% of the annotated genes encode putative roles in stress responses, including the aforementioned Woronin body major protein; a glycoside hydrolase family 17 protein (CAZyme); one gene with a putative link to the HOG1 pathway - peptidase M3A and M3B, thimet/oligopeptidase F (Blyth et al., 2023); three genes encoding oxidoreductase enzymes (oxidoreductase like protein, NAD(P)-binding proteins and FAD/NAD(P) binding proteins) which regulate redox homeostasis and play critical roles in response to oxidative stress and virulence (Kim et al., 2009); a transcription factor C6 like protein (Zn2Cys6), which are multifunctional regulators of many fungal processes including response to osmotic, heat and oxidative stress (Hou et al., 2020); and a gene encoding Rho1 guanine nucleotide exchange factor 3 that is related to cell wall integrity. The glycoside hydrolase family 17 protein is a carbohydrate-active enzyme (CAZyme). These enzymes play a crucial role in pathogen-plant interactions because they are involved in the breakdown of the plant cell wall and can elicit host immune responses. In *Z. tritici,* overexpression of a CAZyme, the glycoside hydrolase *ZtGH45* (Glycosyl hydrolase family 45) leads to a reduced fungal virulence (Rebaque et al., 2025). We speculate that genes within this plGR.4 region could be part of an extended gene cluster that encodes functionally related proteins affecting trait plasticity.

In several cases the genomic intervals harboring multiple plQTL overlap with intervals previously identified in *Z. tritici* QTL mapping studies. In these cases we can draw on knowledge about genes determining variation in the mean traits to identify potential pleiotropic candidate genes for plasticity. The interval on chromosome 11 (GR.23, plGR.18) is one good example. Variation in mean and sensitivity in growth and melanisation traits in the presence of multiple stressors, including fungicide (Lendenmann et al., 2015), temperature (Lendenmann et al., 2016; Stapley et al., 2025), reactive oxygen (Zhong et al., 2021), and salt (Stapley and McDonald, 2023) have consistently mapped to this genomic region, many with LOD >10. This region includes a polyketide synthase (PKS) gene cluster, containing *Pks1* which is involved in DHN melanin biosynthesis, *Zymoseptoria* melanin regulation 1 (*Zmr1*) and 1,3,8-trihydroxynaphthalene reductase (*Thr1*) (Krishnan et al., 2018). In the 3D7x3D1 population, variation in gene expression in *Zmr1* explained variation in colony melanisation *in vitro* and gene expression variation was due to two independent mutations in the 3D1 parent, one being a SNP in the promoter, and the other being a transposable element (TE) in the promoter (Krishnan et al., 2018). There is a known trade-off between melanisation and growth rate, with more melanized strains on average growing more slowly (Krishnan et al., 2018; Lendenmann et al., 2015). Thus variation in *Zmr1* expression can directly affect melanisation rate and indirectly affect growth rate. Mapping of plasticity and mean traits identified associations within this interval, with the highest LOD interval (22.6 for rxnm.grwth.temp) including *Pks1* but not *Zmr1*. There may be some uncertainty in these QTL interval boundaries, as there is a TE in the promoter of *Zmr1* that could create some uncertainty in estimating the 95% confidence intervals for this QTL because we are mapping to the ZT99_3D7 genome, and the presence of a TE in offspring carrying the 3D1 haplotype may result in poor mapping results. Lendenmann et al (2014) previously proposed that one of two nonsynonymous SNPs (amino acid position 884, 1783) within *Pks1* could be responsible for variation in melanisation, favouring the SNP changing proline to threonine (position 1783). Our mapping identified a very small (49 bp), high LOD (25.81) QTL for mean melanisation rate in pH 7.5 within the *Pks1* gene (Table 1) and the peak marker in the ZT3D7 genome (Chr11:658119) was the nonsynonymous SNP that alters the 884th amino acid from a valine to alanine. Thus our mapping results strongly support a role for this SNP in *Pks1* in affecting variation in melanisation rate.

Typically, polyketide synthase genes occur within a BGC together with additional biosynthetic genes such as those encoding cytochrome P450 oxidoreductases, transferases and transcription factors that regulate gene expression within the cluster (Cairns and Meyer, 2017; Jia et al., 2021). Clustering facilitates coordinated gene expression and the BGC can further adapt to environmental stress through horizontal gene transfer, gene loss and rearrangements (Jia et al., 2021; Zeng et al., 2018). One of the enriched GO terms for plasticity, “mycotoxin biosynthesis”, includes genes often arranged in highly plastic, environmentally responsive BGCs (W. Wang et al., 2022). Our mapping results suggest that a polyketide synthase gene cluster could be responsible for the plasticity hotspot plGR.18 on Chr11, where plasticity in response to three different environments maps to a single region. plGR.18 had two BGCs (almost significantly enriched for BGCs p=0.06), we identified a strong candidate SNP within *Pks1*, and this region contains TE copy number variation between the parents - generating genomic rearrangements. Our genetic mapping results support a growing body of research demonstrating that polyketide synthase gene clusters can affect both trait mean and trait plasticity, and strongly support a role for BGCs in the evolution of plasticity.

Another notable genomic region was plGR.17 on chromosome 10. In this case three plasticity traits measuring variation across temperatures (10-18^0^C), salt concentration, and reactive oxygen concentration mapped to this high LOD interval (Table 1). This interval contains *Pbs2*, a well known map kinase kinase (MAPKK). Multiple stressors, including osmotic, heat, cold and pH, activate the HOG1 pathway and Pbs2 phosphorylates this activated Hog1, which in turn regulates expression of many genes (Panadero et al., 2006; Tian et al., 2016). A QTL for cold sensitivity (15-22^0^C) was previously found in this genomic region and *Pbs2* was identified as a likely candidate gene for this trait (Lendenmann et al., 2016). Two corresponding mnQTL (mn.mel.18^0^C, mn.grwth.10^0^C) overlapped the three plQTL with high LOD. In this case, we hypothesize that Pbs2 is acting as a master regulator for both the mean traits and the plasticity traits.

Plasticity in pycnidia grey value and other mean pycnidia traits measured in the 1A5x1E4 population map to a genomic region on Chr5 (GR.10). The plQTL contained 28 genes, including the avirulence factor *AvrStb6* (ZtIPO323_060700). This gene was identified as an avirulence factor following QTL mapping of virulence as a binary trait on Chinese Spring and Drifter in the 1A5x1E4 population, and following GWAS analysis in 103 strains collected in France and one from the UK from fields planted with different cultivars (Zhong et al., 2017). AvrStb6 is a secreted effector protein that triggers an immune response in wheat cultivars carrying the resistance gene *Stb6* (Zhong et al., 2017). We consider it highly plausible that this is an example of trait plasticity due to pleiotropic effects of different alleles of the same gene.

Plasticity in growth rate across variation in H_2_O_2_ and plasticity in melanisation rate across variation in temperature (10-18^0^C) map to an interval on Chr12 which contains the gene *ZtMsr1* encoding a transcription factor controlling a temperature sensitive yeast-hypha morphological transition (Francisco et al., 2023). It also contains a TFIID-18kDa-domain-containing protein, which coordinates transcription initiation. Both of these genes are evolutionarily conserved across eukaryotes. In *Z. tritici, ZtMsr1* was found to be involved in sensing and/or responding to changes in cell wall integrity (CWI) in response to heat stress, however deletion of *ZtMsr1* does not alter the response of cells to oxidative stress (Francisco et al., 2023). The role for the TFIID-18kDa-domain-containing protein within *Z. tritici* is only inferred by homology; across fungi in general no specific role has been shown in oxidative stress or cold temperature stress. Therefore, we propose that *ZtMsr1* is the more plausible candidate gene governing plasticity in growth under H_2_O_2_ stress and melanisation under cold stress.

The highest LOD recorded (31.3) was for a plasticity QTL for rxnm.PyCm2 located on Chr7. This interval contained 28 genes and it overlapped with 19 other QTL (all found within GR.18). The GR.18 region contained five QTL with high LOD ranging from 29.4-31.3, and included nine mnQTL (mnQTL.15) and 11 plQTL (plGR.12, and plGR.13). plGR.12 contained two plQTL for melanisation rate measured across variation in temperature and plGR.13 contained nine *in planta* plasticity trait QTL (Table 2). GR.18 (and the plGR within) contains the gene encoding Avr3D1, a small secreted protein that triggers quantitative resistance in cultivars with the Stb7 resistance gene (Meile et al., 2018). The region also contains a gene encoding a common fungal extracellular membrane (CFEM) domain-containing protein and a glycoside hydrolase family 36 protein (CAZyme). The CFEM domain-containing proteins are fungal-specific, cysteine-rich effectors crucial for virulence (Iliopoulos et al., 2026), maintaining cell wall integrity (e.g. *Fusarium verticillioides* (Li et al., 2025)) and suppressing plant immunity (e.g. *Verticillium dahliae* (D. Wang et al., 2022)). The role of CAZymes in fungal virulence was discussed earlier. We speculate that plGR.12, and plGR.13 are examples of a chromosome segment that contains several tightly linked genes that independently affect plasticity of several traits associated with pathogen-plant interactions. This could generate complex fitness trade-offs that would predict a rapid evolution of the affected genes in these segments.

Next we consider the few cases where the plQTL and the corresponding mnQTL mapped to different regions of the genome, thus supporting the epistatic model where genes affecting plasticity evolve independently from genes affecting the mean trait. These cases included three plasticity traits in the 1A5x1E4 population: RDPI in growth rate and melanisation rate across variation in reactive oxygen concentration, and RDPI in growth rate across variation in temperature (10-18^0^C); and one trait in the 3D7x3D1 population: RDPI in melanisation rate across temperature variation (18-27^0^C). All of these are RDPI plasticity measurements that have relatively low LOD scores (ranging from 4.52-7.68), thus these QTL have relatively small effect sizes. The correlation coefficients between these plasticity traits and their means were significantly lower than the pairs of plasticity-mean traits with overlapping QTL, supporting the interpretation that these traits are controlled by independent loci. Within the epistasis model, variation in plasticity is driven by environmentally sensitive gene regulatory networks (GRNs), composed of a regulator (like a transcription factor) and its target locus or signaling pathway (Gulisija and Plotkin, 2017; Scheiner, 1993). One modelling study that explored how selection on plasticity can drive the evolution of recombination between an environmentally responsive regulator (plasticity locus) and the target locus controlling the mean, or lead to the tight clustering of these genes (as we proposed for GR.18), has shown that the outcome is dependent on the frequency of the environmental variation, epistasis and the number of interacting genes (Gulisija and Plotkin, 2017). Further studies focused on the distinct genomic regions harbouring plGR and mnQTL identified in this study, could provide novel insights into epistasis and the evolution of plasticity. Fungal models like *Z. tritici* offer many advantages compared to plants and animals for understanding the molecular mechanisms driving the evolution of plasticity, including both sexual and asexual reproduction, small genomes, easily measured phenotypes across different environments, amenability to experimental evolution, and access to a wide range of molecular tools that can readily identify causal mutations affecting trait plasticity.

GO enrichment analyses of plQTL identified two enriched GO annotations; mycotoxin biosynthetic process and serine-type peptidase activity. When considering only genomic regions containing plQTL explaining trait variation across different environments, three GO annotations were enriched; DNA-directed DNA polymerase activity (DNA-PA), proteasome-activating activity (PAA) and serine-type peptidase activity (SPA). DNA-PA could influence phenotypic plasticity because it plays a central role in maintaining epigenetic marks and DNA repair. Epigenetics have been found to play a role in phenotypic plasticity across environmental variation. In *Neurospora crassa* epigenetic modifications were more important to plasticity in growth compared to variation in the mean growth (Kronholm et al., 2016). Genes encoding PAA could also play a role in phenotypic plasticity in fungi. Proteases were shown to interact with Hsp90 and Hsf1 in maintaining global protein homeostasis during stress in multiple *Candida* species as well as regulating fungal morphogenesis (Hossain et al., 2020). *SPA* genes encode essential, highly diverse, and frequently secreted enzymes that play key roles in nutrient acquisition, environmental adaptation, and pathogenicity (Muszewska et al., 2017). In *Botrytis cinerea* a strong heat priming effect, in which exposure to moderately high temperatures greatly improves a strain’s ability to cope with subsequent, potentially lethal temperature conditions, was linked to a group of priming-induced serine-type peptidases (Zhang et al., 2023). The results of these studies, combined with our GO enrichment analysis, support our hypothesis that these genes are also plausible candidates for affecting fungal phenotypic plasticity.

### Are phenotypic correlations indicative of shared genetic control?

There is a general assumption that phenotypic correlations reflect underlying genetic correlations – which are indicative of shared genetic control (pleiotropy) or tight linkage between different genes. However environmental factors can also drive these trait correlations. We found that most plQTL overlapping with mnQTL also had high mean absolute correlation coefficients (mnACC) between plasticity and mean phenotypic traits (mnACC=0.65) (Supplementary Table S5a). However, there were two pairs of traits where there was QTL overlap (i.e. traits were genetically correlated) but the phenotypic traits were not correlated. For example, in the 1A5x1E4 population the QTL for rxnm.mel.Temp overlaps with mn.mel.18^0^C and mn.mel.15^0^C. However, rxnm.mel.Temp is not correlated with mn.mel.18^0^C (correlation coefficient (CC) =0.07), instead it is strongly negatively correlated with mn.mel.15^0^C (CC=-0.85) and positively correlated with mn.mel.22^0^C. This is likely due to the opposing environmental effects and pleiotropy which is captured in the quadratic reaction norm. A similar result was observed in the 3D7x3D1 population: rxnm.grwth.KCl is not strongly correlated with mn.grwth.KCl (CC=0.17), but is negatively correlated with mn.grwth.18^0^C (CC=-0.71). We can also consider the reciprocal of this argument - do traits mapping to distinct genomic regions have low phenotypic correlation coefficients? If we consider the four plasticity traits that had plQTL that never overlapped with the corresponding mnQTL, the phenotypic mnACCs were significantly lower (mnACC=0.36) than the mnACC between traits with overlapping QTL (mnACC=0.65). Therefore we can say the correlations were lower, on average, for these four traits, however most correlations were significant (15/18), and in one case (lrdpi.mel.H2O2) the correlation between the plasticity and mean trait was as high as -0.61 (Supplementary Table S5b). Overall, our results suggest that in most cases a phenotypic correlation between trait mean and trait plasticity does reflect shared genetic control, but significant phenotypic correlations can also exist between traits that are controlled by distinct genetic loci. Thus, we must be cautious when drawing conclusions about the genetic control of correlated phenotypic traits.

## Conclusions

In this study we simultaneously mapped trait plasticity and trait means for both *in vitro* and *in planta* traits in experimental populations of an important fungal plant pathogen. The traits were measured across a broad array of both abiotic and biotic environments. We identified multiple significant QTL and GWAS associations, providing unprecedented insights into the genetic architecture of trait plasticity in a plant pathogen. We found that in most cases the QTL for trait plasticity and the corresponding trait mean mapped to the same genomic regions, suggesting that plasticity is controlled by pleiotropy and/or tightly linked genes. We also found hotspots in the genome where plasticity measured across variation in different environments mapped to the same region, suggesting shared genetic control. These hotspots contained plausible master regulators of plasticity (e.g. *Hex-1*, *Hsf1*) and biosynthetic gene clusters (e.g. *Pks* gene clusters). This study provides important insights into the genetic architecture and evolution of plasticity in pathogenic fungi.

## Acknowledgements

This work was inspired by a conversation with Anna-Liisa Laine at University of Zurich in January 2020. We thank the principal data collectors and their teams for phenotypic data collection: Mark Lendenmann, Ethan Stewart, Ziming Zhong, and Anik Dutta. Special thanks go to Marcello Zala for his constant technical support. Funding was provided by Swiss National Science Foundation (31003A_134755, 31003A_155955) and the Swiss Federal Office for Agriculture (BLW) in the framework of NAP-PGREL (National Plan of Action for the Conservation and Sustainable Utilization of Plant Genetic Resources) Project Nr. 627000640. Genomic data analysis was supported by the Genetic Diversity Centre (GDC).

## Author Contributions

JS: formal analysis, data curation, writing—original draft, writing—review & editing, resources, and data curation. BAM: conceptualization, methodology, writing—review & editing, supervision, project administration, and funding acquisition.

## Data availability

Data are available at https://www.research-collection.ethz.ch/ handle/DOI:XXX. Metadata, code, and details of the analysis are available at https://github.com/jessstapley/QTL-mapping-Z.-tritici.

## Supplementary Tables

**Table S1.** Description of 145 *Zymoseptoria tritici* strains used for GWAS analyses in this study, including their sampling location, year and the NCBI SRR Run ID for the whole genome sequence datasets.

**Table S2.** List of traits mapped using QTL or GWAS analyses. For *in vitro* traits, growth rate and melanisation rate were measured across variation in an environment. Plasticity was calculated from measurements taken in two or more environments [e.g. benign-stressful (PDA-Salt, PDA-Fungicide); variations in temperature 10C, 15C, 18C, 22C, 27C]. Trait (PDA-Salt, PDA-Fungicide); variations in temperature 10C, 15C, 18C, 22C, 27C]. Trait mean was calculated in a single environment [e.g. in the benign control (PDA), in the presence of a Fungicide, in low pH (4.5)]. For *in planta* traits, Ethan Stewart measured pycnidia area, pycnidia count, pycnidia density, pycnidia grey value, and percent leaf area covered by lesions (PLACL) in two cultivars (Runal, Titlis); Anik Dutta measured reproduction traits (pycnidia-based) and virulence (PLACL-based) across eight cultivars (ArinaLr34, Chinese Spring, Drifter, Gene, Greina, Runal, Titlis and Toronit).

**Table S3.** The number of significant associations (using either QTL or GWAS analyses) for all traits for each experimental population (1A5x1E4, 3D7x3D1, global) and the number of experimental populations (n.populations) with at least one significant association.

**Table S4a.** Summary of the genomic regions (GRs) containing significant associations based on QTL or GWAS analyses for all traits. For each GR the table lists the chromosome (Chr), the region start and stop positions and length in base pairs, the experimental population used for mapping (Pop), the experimenter’s initials (Experimenter), the number of traits with significant QTL (n.QTL traits), the number of traits with significant GWAS associations (n.GWAS traits), the number of significant GWAS SNPs (n.GWAS SNPs), the number of different environments in which the traits were measured (n.Env), the number of biosynthetic gene clusters (n.BGC) in the region and a list of all traits associated with the region.

**Table S4b.** Summary of the genomic regions (GRs) containing significant associations based on QTL or GWAS analyses for plasticity traits.

**Table S4c.** Summary of the genomic regions (GRs) containing significant associations based on QTL or GWAS analyses for mean traits.

**Table S5a.** Details of QTL for plasticity (plQTL) traits that overlap with their corresponding mean trait QTL (mnQTL), including the QTL coordinates (chromosome, start, stop), experimental populations used in the mapping (Pop), the length of overlap in base pairs, the percentage of the plQTL that is overlapping (% plQTL), the percentage of the mnQTL that is overlapping (% mnQTL), the phenotypic correlation coefficient between the plasticity trait and the mean trait (correlation), and significance of the correlation (p-value). A plasticity trait can have multiple plasticity QTL and each plasticity trait has multiple mean corresponding traits, thus a plQTL can overlap with all or some of the corresponding mnQTL (QTL.overlap). Alternatively one plQTL can have no overlap with any corresponding mnQTL, while other plQTL for the same trait can overlap a mnQTL (no.overlap). There are cases where all plQTL for a single trait never overlap a corresponding mn.QTL.

**Table S5b.** QTL for plasticity traits that do not overlap with their corresponding mean trait QTL.

**Table S5c.** GWAS SNPs for plasticity traits that do not overlap with their corresponding mean trait GWAS SNPs.

**Table S6.** Results from GO enrichment analyses.

**Table S7.** Annotated candidate genes located within the most significant plasticity trait QTL described in the main text. The most plausible candidate genes to explain the observed phenotype are colored in red.

**Table S8.** A list of significant GWAS SNPs with details of the corresponding genomic regions (GR), the trait type (pl: plasticity, mn:mean), *in vitro* or *in planta*, abbreviated trait name, SNP information and gene position in the IPO323 genome, p-value and negative Log p-value (nLpv), minor allele frequency (MAF) of SNP, the estimated effect, information about the gene where the SNP is located (gene name, position and annotation) or intergenic if the SNP does not overlap with an annotated gene.

## Supplementary Figure legends

**Figure S1.** Density histogram of the absolute correlation coefficient (ACC) between pairs of phenotypic traits. Orange shows ACC between a plasticity trait and the corresponding mean, grey shows ACC between all pairs of traits. Lines indicate mean absolute correlation coefficients.

**Figure S2**. Heatmaps of absolute correlation coefficients between plasticity traits for experimental populations: a) 1A5x1E4, b) 3D7x3D1, c) global *in planta* traits, and d) global *in vitro* traits.

**Figure S3**. SNP heritability (h^2^) for plasticity traits and mean traits measured *in vitro* and *in planta*. Experimental populations include the two QTL crosses (1a = 1A5 x 1E4; 3d = 3D7 x 3D1) and the GWAS global population. Letters above bars show grouping according to least significant difference (LSD).

**Figure S4**. Relationship between the number of significant associations and chromosome length for each population (1A5x1E4, 3D1x3D7, global). Points are jittered (random horizontal noise) to reveal overlapping points.

**Figure S5.** Density histograms of the percentage overlap between QTL intervals. a) overlap between a plQTL and the corresponding mnQTL, b) overlap between all pairwise plQTL, c) overlap between all pairwise mnQTL, and d) overlap between all pairwise QTL.

## Supplementary Material

Tables are Comma separated text (csv) file.

**Table SM1.** Summary of the linkage map length and estimated genome coverage.

**Table SM2.** Pairwise phenotypic correlations for *in vitro* traits measured in the 1A5x1E4 population.

**Table SM3.** Pairwise phenotypic correlations for *in vitro* traits measured in the 3D7x3D1 population.

**Table SM4.** Pairwise phenotypic correlations for *in vitro* traits measured in the global population.

**Table SM5.** Pairwise phenotypic correlations for *in planta* traits measured in the global population.

**Table SM6.** Phenotypic data used in QTL mapping of the 1A5x1E4 population.

**Table SM7.** Phenotypic data used in QTL mapping of the 3D7x3D1 population.

**Table SM8.** Phenotypic data used in GWAS mapping of *in vitro* traits measured in the global population.

**Table SM9.** Phenotypic data used in GWAS mapping of *in planta* traits measured in the global population.

**Table SM10.** Comparison of the *Hex-1* gene model across *Z. tritici* strains. The reference sequence is from IPO323 and the four QTL cross parents are 1A4, 1E4, 3D7, 3D1.

**Table SM11.** Comparison of the *Hsf* gene model across *Z. tritici* strains. The reference sequence is from IPO323 and the four QTL cross parents are 1A4, 1E4, 3D7, 3D1.

