## Supplementary figures and images for "Genetic mapping of trait plasticity in a plant pathogenic fungus reveals genetic architecture and candidate genes for plasticity"

### Figure S1

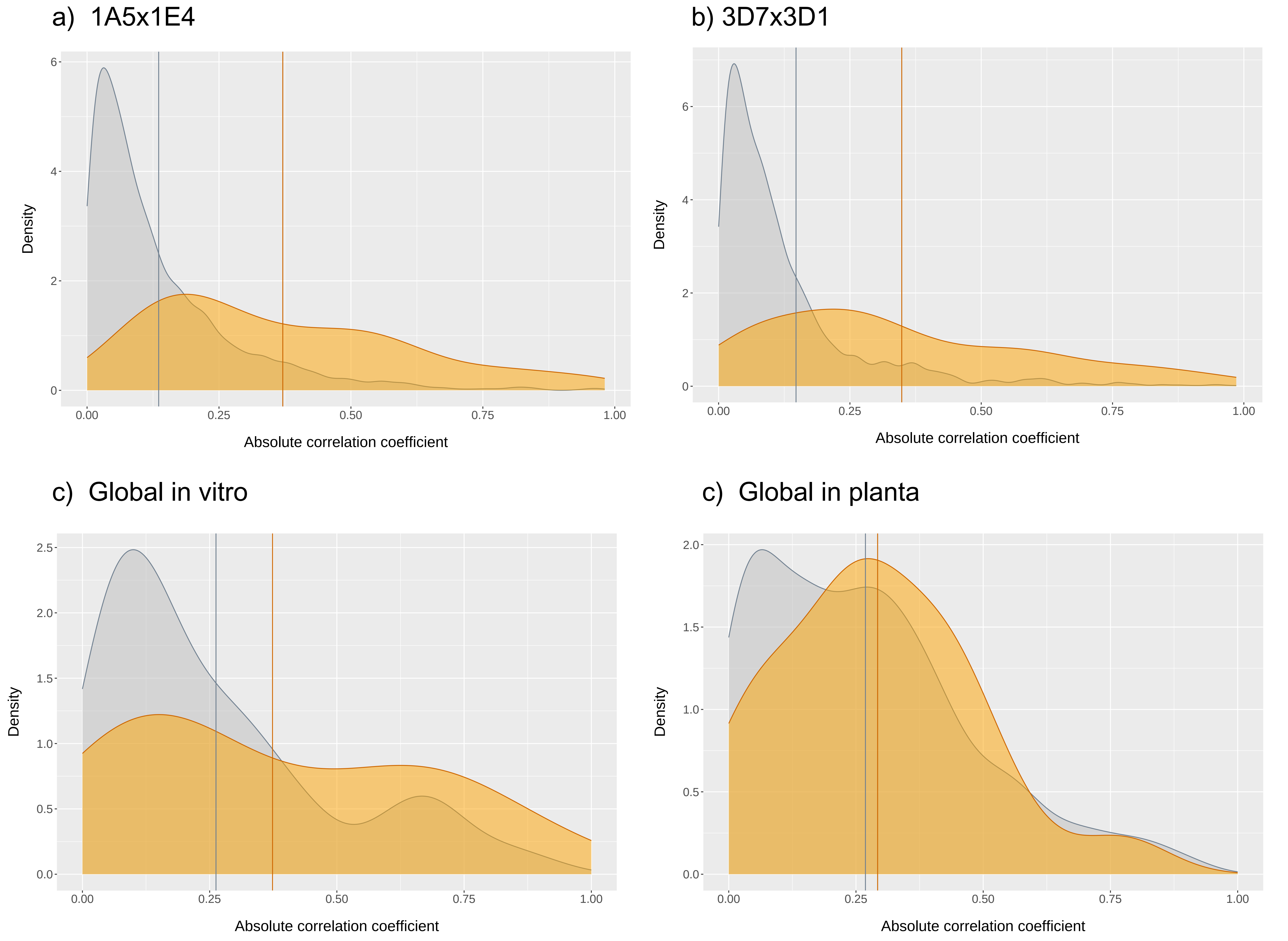

### Figure S2

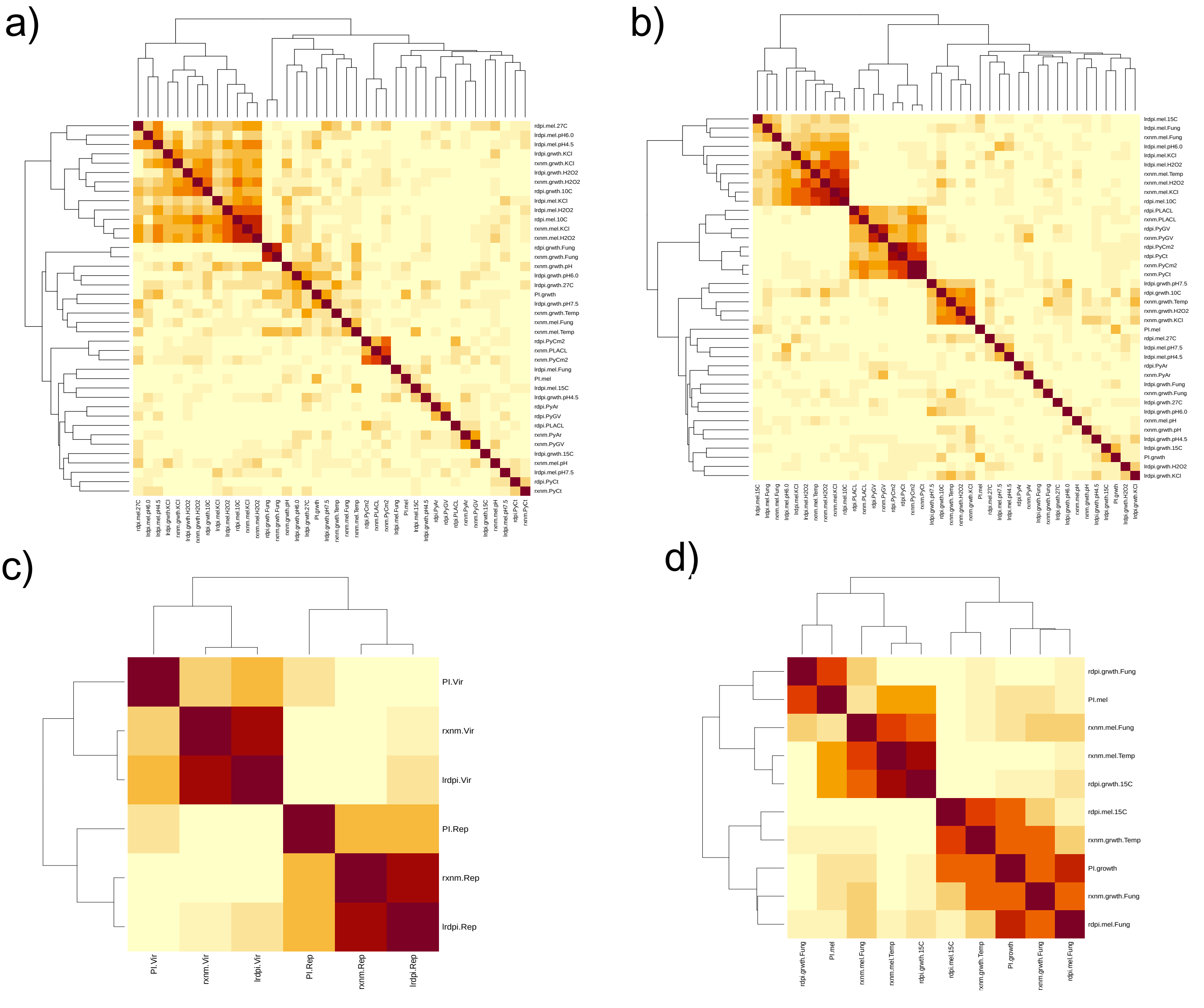

### Figure S3

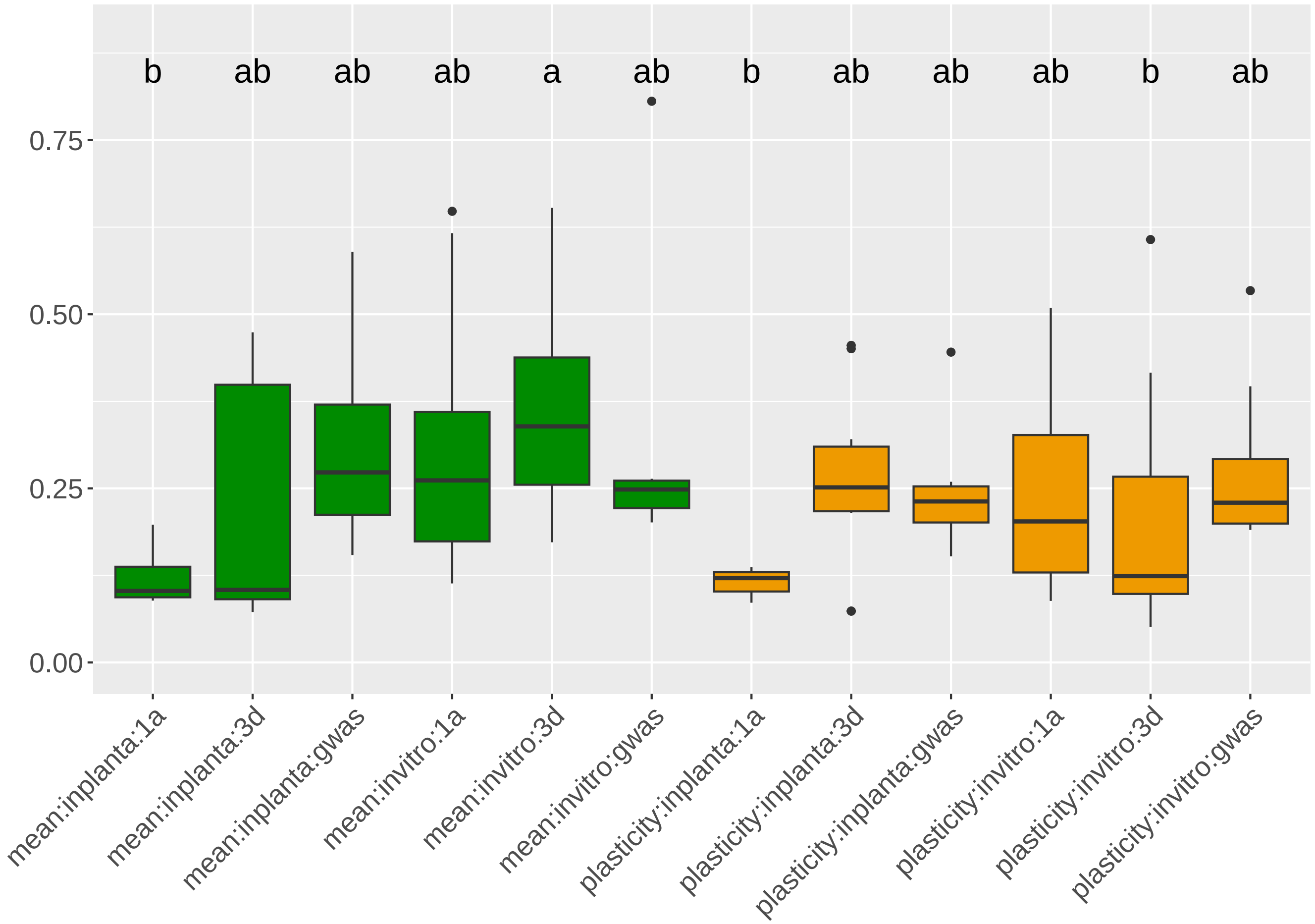

### Figure S4

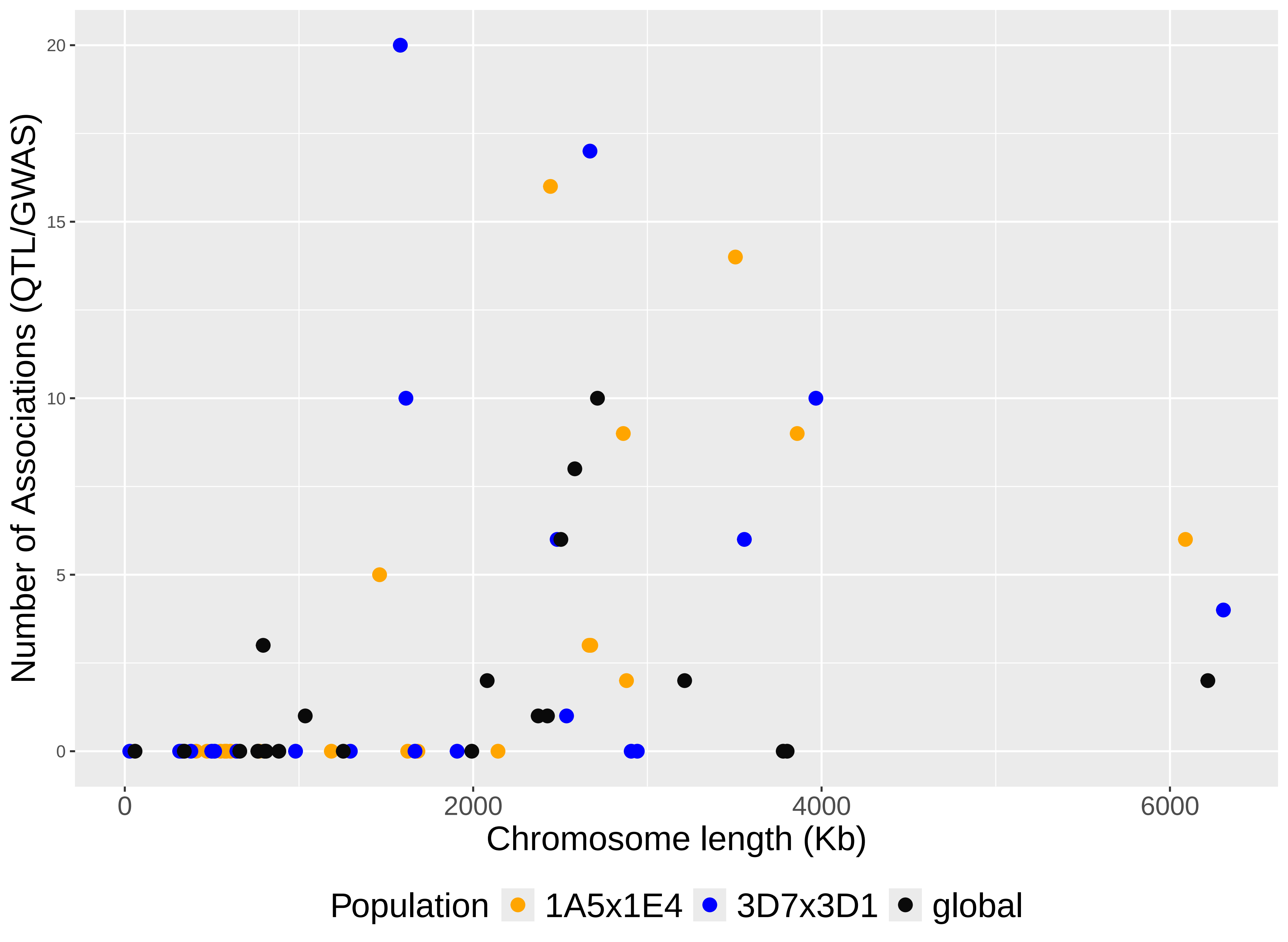

### Figure S5

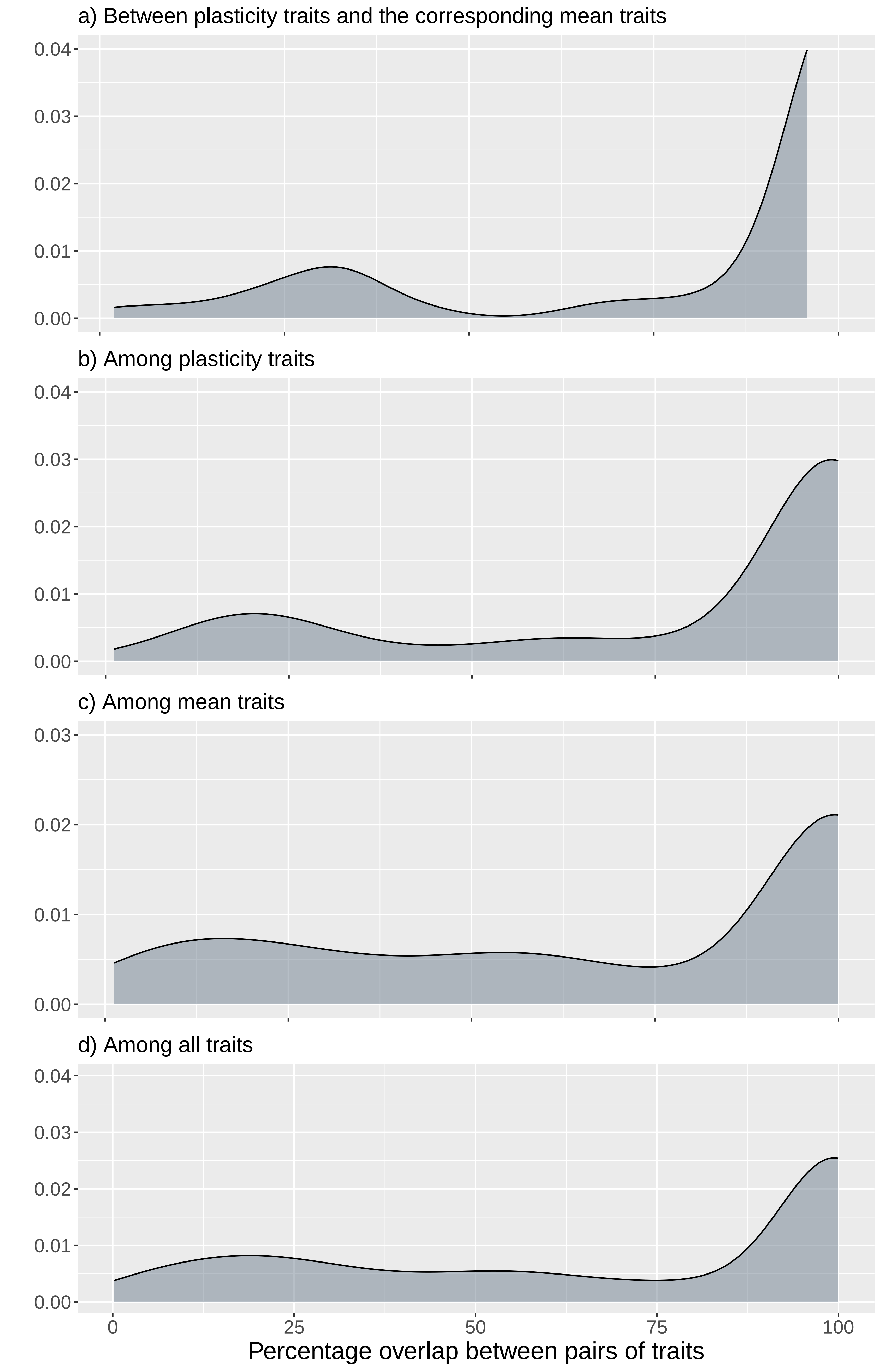
